# Reconstruct a Connectome of Single Neurons in Mouse Brains by Cross-Validating Multi-Scale Multi-Modality Data

**DOI:** 10.1101/2024.10.01.616182

**Authors:** Feng Xiong, Lijuan Liu, Hanchuan Peng

**Author notes:** Equal contribution (Feng Xiong, Hanchuan Peng). Correspondence: Hanchuan Peng.

## Abstract

Brain networks, often referred to as connectomes, are crucial to findings in brain science and have inspired the development of numerous technologies at macro-, meso-, and micro-scales. With the growing prevalence of single-cell analysis, there is an urgency to generate connectomes of individual neurons that project to brain-wide regions. This work reports a scalable approach to build neuron connection networks of mouse brains by mapping whole-brain single-neuron connectivity using two complementary methods at sub-neuronal levels. We first generated an arbor-net by partitioning and probabilistically pairing dendritic and axonal arbors of 20,247 neurons registered to the Allen Brain Atlas. We also produced a bouton-net based on 2.57 million putative boutons detected along single axons of 1,877 fully reconstructed neurons and probabilistic pairing of these full-morphology datasets. Our cross-validation of both the arbor-net and bouton-net showed statistical consistency in the spatially and anatomically modular distributions of neuronal connections, which also corresponded to functional modules of the mouse brain. Our single-neuron connections were also validated by two existing independent connectomes of coarser resolution based on viral tracing of neuron populations and barcoding-and-sequencing, as well as an independent synaptome containing the relative density distribution of synapses. We further found that single neuronal connections correlated more strongly with gene co-expression at both the brain region and single-cell levels than the previous full-brain mesoscale connectome of a mouse brain. Our findings allow the assembly of a new whole-brain scale single-neuron resolution connectome for all brain regions, called SEU-net. We studied the properties of our connectomes in comparison with other potential brain architectures and found a series of non-random subnetwork patterns in the form of consistent triad motifs. Overall, our data indicate a rich granularity and strong modular diversity in the brain networks of mice.

## Introduction

A connectome (Sporns, et al, 2005; Lichtman, et al, 2008; DeFelipe, 2010; Van Essen, et al, 2012; Van Essen, et al, 2013; Bargmann and Marder, 2013; Sporns, 2013; Oh, et al, 2014), is a complete representation of neuronal connections among involved neurons, neuronal populations, and brain regions. Connectomes have been intensively studied at various scales, including the macro-scale (e.g., Van Essen, et al, 2012; Van Essen, et al, 2013), mesoscale (e.g., Oh, et al, 2014; Hunnicutt, et al, 2014), and microscale (e.g., Yin, et al, 2020; Turner, et al, 2022; Shapson-Coe, et al, 2024). These connections can be derived from structural neuronal data, such as neuronal morphology (Liu, Yun, et al, 2023), functional neuronal data, like physiological recordings (Franconville, et al, 2018), or genetic approaches such as DNA barcoding (Zador, et al, 2012).

Clearly, any partial representation of these connections compromises the accuracy and significance of such connectomes. Practically, different resolutions and approaches for constructing connectomes involve various compromises. For example, while electron microscopy (EM) can precisely identify individual synaptic contacts, it often works with limited volumes of brain tissue. Consequently, EM faces challenges in fully capturing the 3-D morphology and long-range projection information of brain-scale projecting neurons needed to define neuronal identities uniquely. Without this information, a microscale connectome constructed using EM would be insufficient for understanding the brain-scale wiring patterns and neuron distribution (Rees, et al, 2017).

With the increasing availability of single-cell neuron morphometry studies (Winnubst, et al, 2019; Peng, et al, 2021; Gao, et al, 2022; Liu, Yun, et al, 2023; Liu, Jiang, et al, 2023; Qiu, et al, 2024), it is now desirable to study how to infer brain networks with both local and long-range projections based on individual neurons (e.g. Qian, et al, 2024). Progress may be made using either a single brain or multiple brains, the latter requiring multiplexing with 3-D brain registration techniques but also capable of incorporating a tremendous amount of multimodal information (e.g., Qu, et al, 2022). Compared to an established mesoscale projectome that relies on neurite bundles (Oh, et al, 2014), the increased spatial resolution provided by single neuronal projections is a remarkable advantage, particularly because it enables visualizing the long-range organization of brain networks, potentially even at the whole-brain scale, with neuron or sub-neuronal precision. These long-range neuronal projection patterns also enhance cell typing (Liu, Yun, et al, 2023; Xiong, et al, 2024), which is an orthogonal effort identifying modules in brain anatomy.

This work reports a scalable approach to build neuron connection networks in mouse brains by mapping whole-brain single-neuron connectivity using two complementary methods at sub-neuronal levels. We created an arbor-net and a bouton-net by pairing multiple independently generated axonal and dendritic features, all registered to the Allen Brain Atlas, both spatially and probabilistically. We cross-validated them, highlighting their statistical and functional consistency in the spatial and anatomical modular distributions of neuronal connections. We also cross-validated our connectomes with existing independent connectomes of coarser resolution based on viral tracing of neuron populations and barcoding-and-sequencing, and an independent synaptome with brain-wide distribution of synapses. We additionally discovered that single neuronal connections correlated more strongly with gene co-expression at both the brain region and single-cell levels than the previous full-brain mesoscale connectome of a mouse brain. We compared our connectomes with other potential brain architectures and identified a series of non-random motif-networks. Overall, our data reveal rich granularity and strong modular diversity in the brain networks of mice that have not been quantitatively characterized before. Our work also highlights a traditionally used approach, Peters’ rule (Braitenberg and Schüz, 2013), alone is not sufficient; it should be conjugated with other evidence and can be conveniently enhanced by using statistics shown in this study.

## Results

### A framework to build connectomes of single-neurons cross-verifiable by gene co-expression

In the following, to avoid confusion, we use the term “connectome” to refer to a connection network for a specific set of neurons. In particular, we built connectomes based on the 3-D full morphologies of 1,877 neurons (called “F1877” for simplicity) and the dendritic morphologies of 18,370 neurons (called ‘D18370’), which were reconstructed using a recently developed neuroinformatics platform as discussed in a related article (Liu, Jiang, et al, 2023; **Methods**). For the fully reconstructed neurons in F1877, we also generated 2.57 million Predicted Boutons (PBs) along their axonal arbors (**Methods**). We registered spatially all these neuron morphologies and PBs to the Allen Common Coordinate Framework (CCF) version 3 atlas (Wang, et al, 2020) using the mBrainAligner tool (Qu, et al, 2022). Within the standard atlas space, we identified and analyzed the spatial adjacency of each axon-dendrite pair among these neurons to estimate strength scores of potential synaptic connections (**Figure 1A**).

**Figure 1.**
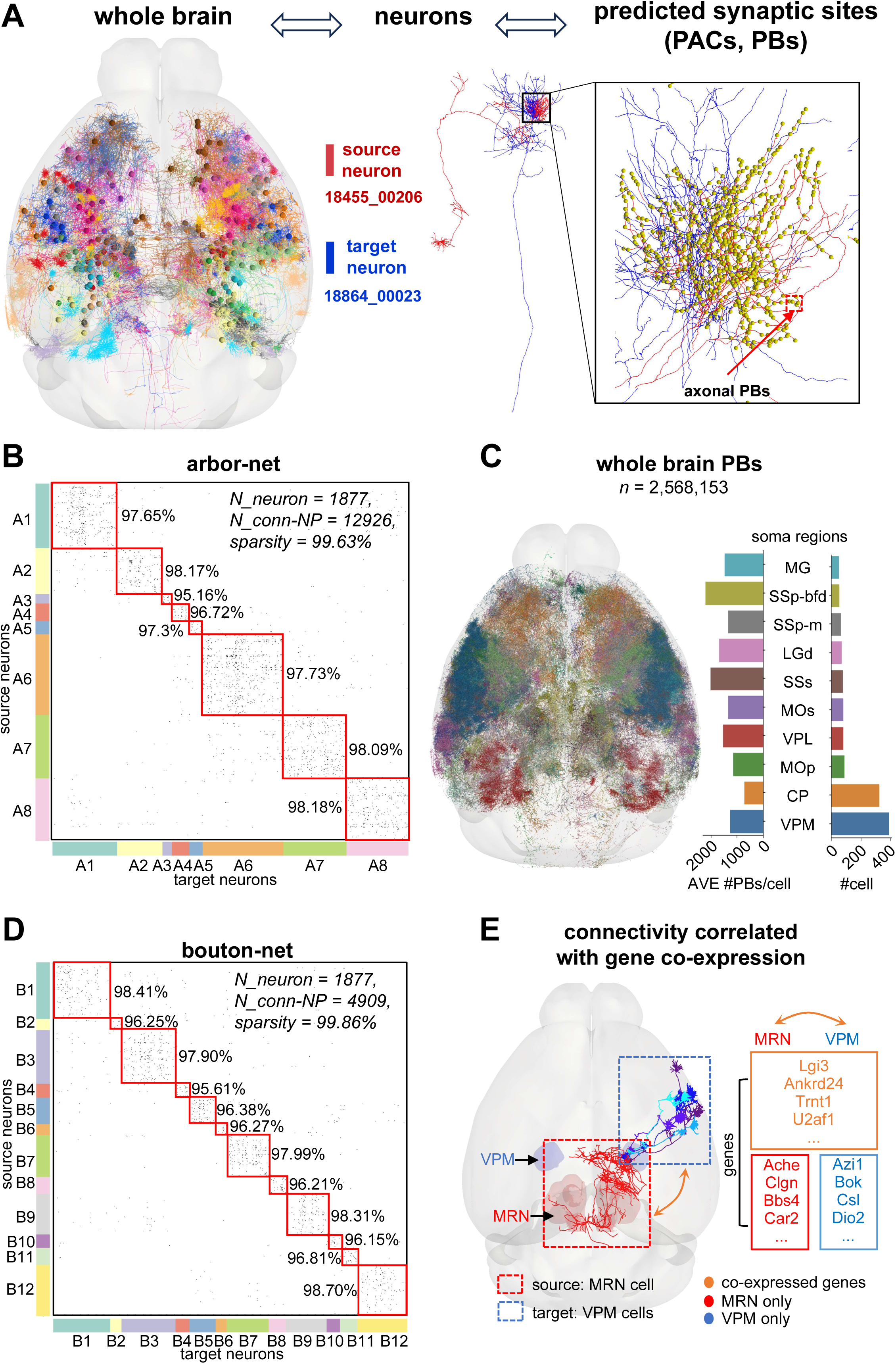
Overall scheme to build the single-neuron level connectomes cross-validated using multi-modality datasets such as gene co-expression. **A**. Overview of multi-level connection networks of single-neurons. Left: a set of full neuron reconstructions in whole mouse brain; Middle: axonal arbors (red) spatially overlap with dendritic arbors (blue); Right: predicted axonal synaptic sites (yellow) spatially approach dendritic arbors. **B.** Arbor-net of 12,926 connections within 1,877 neurons. Vertical axis: source neurons. Horizontal axis: target neurons. Values in matrix: connections (1, black) and non-connections (0, white) derived from PACs. Colors along the axes correspond to A1-A8 modules. Red boxes and percentages: detected modules and sparsity. **C.** Spatial distribution of predicted axonal bouton sites in whole mouse brain. Bar-charts: average counts of PBs per cell in main brain regions (left); number of cells in main brain regions (right). See **Supplementary Table 1** for abbreviations of all brain-region names. **D.** Bouton-net of 4,909 connections within 1,877 neurons. Values in matrix: connections (1, black) and non-connections (0, white) derived from PBs. Colors along the axes correspond to B1-B12 modules. **E.** Whole-brain region connectivity cross-validated by gene co-expression patterns. Left: a group of single-neuron level connections from MRN (red) to VPM (blue). Right: co-expressed genes (yellow) between two brain regions.

We developed two methods to define such connection scores. First, we calculated the shortest Euclidean distance between neurons’ axon arbors and the nearest dendritic morphologies. When the distance was smaller than 5μm, which is the sum of typical length of dendritic spines and the size of axon boutons as reported in literatures (Iascone, et al, 2020; Jiang, et al, 2022), we called the respective axon-dendrite location a Potential Arbor Contact (PAC). We found 427,206 PACs among the fully reconstructed neurons in F1877. As there are typically multiple PACs between a pair of neurons, we then defined the strength score as a scaled function aggregating the number of PACs as well as the spatial adjacency associated with them (**Methods**). This method allowed us to generate an arbor-level connectome, called “arbor-net”, for neurons in F1877 based on their spatial proximity and the receptive profiles of their axonal and dendritic arbors, in terms of 12,926 pairs of potentially connected neurons (**Figure 1B**; **Supplementary Figure 1**). A higher connection strength indicates greater spatial closeness between neurons. Overall, the arbor-net is highly sparse (99.63%). We clustered these neurons into 8 groups, A1 ~ A8 (**Supplementary Figure 2**), using a graph partition algorithm (**Methods**). Within each cluster, neurons have more PACs among themselves and thus appear less sparse than the entire dataset (**Figure 1B**). In another word, when we sorted the arbor-net based on these 8 clusters, we found that the module density within each cluster was higher than that between clusters (**Figure 1B**). The brain region-specific relationships of these clusters will be discussed below.

Second, using spatially registered PBs (**Figure 1C**), we generated a bouton-connectome, called bouton-net, as an alternative to the arbor-net. The key difference was that instead of using PACs, we posited that potential synaptic contacts could occur only at the locations of the predicted axonal boutons. In this bouton-net, we identified 4,909 pairs of potentially connected neurons in F1877 (**Figure 1D**; **Supplementary Figure 3**), with an overall sparsity of 99.86%. Employing the same graph partition algorithm, we clustered the neurons into 12 modules, B1 ~ B12 (**Supplementary Figure 4**), based on which we sorted this bouton-net and found that the module density within each cluster was also higher than that between clusters (**Figure 1D**), similar to the case of coherent modules found in the arbor-net.

Of note, as these two connectomes were produced following essentially different approaches, we further cross-validated them to understand their modules, as shown in the subsequent results (**Figures 2 and 3**). In addition, to ascertain whether these two connectomes might have over- or under-estimated probable neuronal connections, we sought independent evidence. Currently, there is a lack of whole-brain synaptic data derived from electron microscopy, as well as a comprehensive record of the simultaneous firing patterns of upstream-downstream neuron pairs at single-cell resolution. Given these challenges in using synaptic and physiological data, except those we show later in our comparison, this study utilizes gene co-expression data across whole-brain regions to cross-validate potential connectivity patterns in the two connectomes (**Figure 1E**). For instance, we correlated the *in situ* hybridization gene expression data of both VPM (ventral posteromedial nucleus of the thalamus; see **Supplementary Table 1** for all abbreviations of brain regions’ names) and MRN neurons to identify genes that are either shared or uniquely expressed in these populations (**Figure 1E**). A global connection model, encompassing all possible pairs of neurons and their shared genes, provides hints as to whether some connections might be more probable than others.

**Figure 2.**
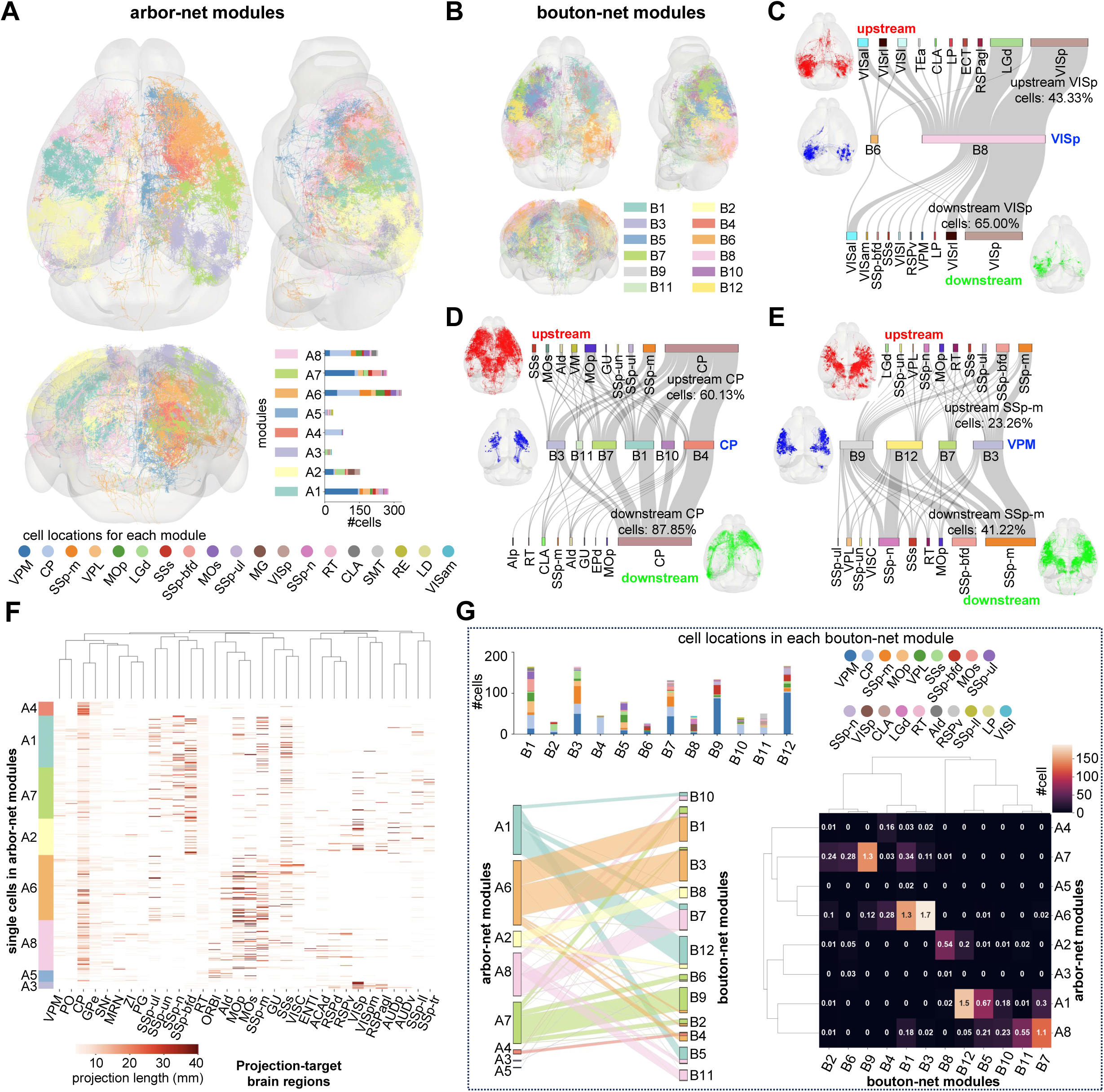
Modules in the arbor-net and their matching modules in the bouton-net. **A-B.** Tri-view visualization of neuron reconstructions in arbor-net and bouton-net modules. Colors indicate the corresponding modules. Bar-chart (bottom right): number of neurons in each module, and their soma locations are indicated by colors. **C-E.** Wiring diagrams for neurons (in three representative regions VISp, CP, and VPM) involving upstream and downstream connections from bouton-net. Connections were curated by expert to prevent false positives, and modules of less than 5% total cells for each brain region were not shown so the predominant patterns are better visualized. Top bars: percentages of upstream connections from various source brain regions; Middle bars: percentages of cells between various bouton-net modules; Bottom bars: percentages of downstream connections targeting to various brain regions. Lengths of bars represent percentages. Aside the bars: 3D-visualization of neuron reconstructions respectively. **F.** Axonal projection of single neurons in arbor-net modules. Vertical axis: source cells, grouped by arbor-net modules. Horizontal axis: axonal projection-target brain regions. Color bar: axonal lengths of cells in target regions. **G.** Comparison of arbor-net modules and bouton-net modules. Bar-chart (top): number of neurons in each bouton-net module, and their soma locations are indicated by colors. Bottom-left: parallel set visualization of neurons. Cells are first aligned according to A1-A8 and B1-B11, and then linked between arbor-net modules and bouton-net modules. Bottom-right: heatmap of shared neurons by arbor-net modules (vertical axis) and bouton-net modules (horizontal axis). Color bar: number of cells.

**Figure 3.**
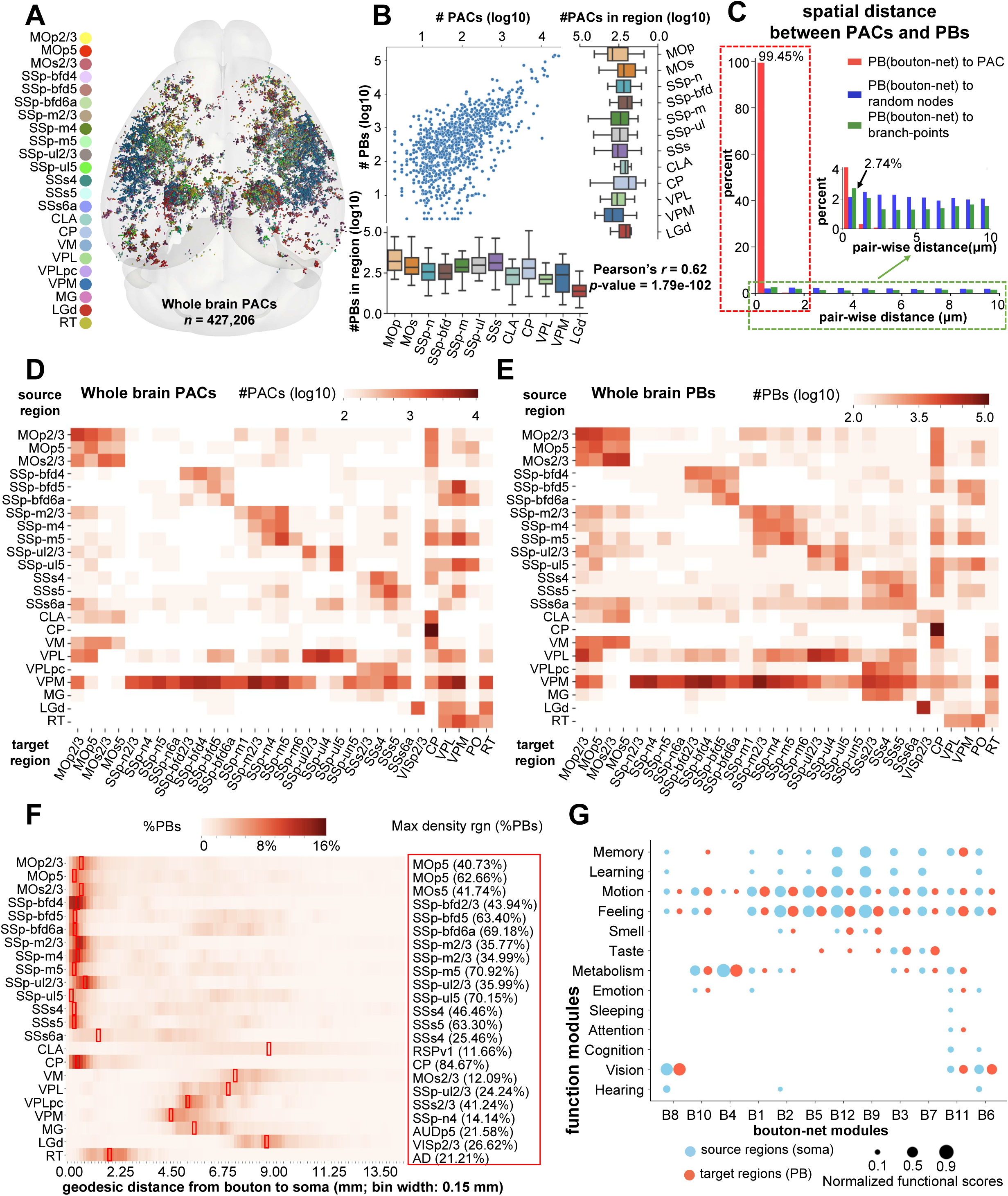
Spatial distributions and co-occurrence of PACs, PBs, and the related function modules. **A.** Spatial distribution of PACs in whole mouse brain. Colors of PACs indicate their soma locations. **B.** Correlation between anatomical distribution between PBs and PACs across whole mouse brain. Scatter plot (top-left) shows the counts of PACs and PBs that are from same source regions and located in the same target regions. Bar-chart (top right): the number of PACs from various brain regions (vertical axis: source brain regions). Bar-chart (bottom left): the number of PBs from various brain regions (horizontal axis: source brain regions). **C.** Euclidean distance between PBs and PACs (red: from PB site to its nearest PAC site; blue: from PB site to its nearest random axonal reconstruction site; green: from PB site to its nearest branching-points). Red box: the major pattern of spatial distance between PBs and PACs. Green box and inset plot indicate the major spatial distribution pattern between PBs and branching-points. **D.** Heatmap of anatomical PACs distribution in whole mouse brain. Horizontal axis: target brain regions of PACs. Vertical axis: source brain regions of PACs. Color bar: the number of PACs (log10-transformed). **E.** Heatmap of anatomical PBs distribution in whole mouse brain. Horizontal axis: target brain regions of PBs. Vertical axis: source brain regions of PBs. Color bar: the number of PBs (log10-transformed). **F.** Predicted bouton sites along axonal reconstruction. Vertical axis (left): source brain regions of PBs. Vertical axis (right): target brain regions of most boutons from corresponding source brain regions, and small boxes on the heatmap also indicate these regions. Horizontal axis: geodesic distance from bouton site to soma location (bin width: 0.15 mm). Color bar: percentages of PBs in each bin. **G.** Relationship between bouton-net modules and brain function modules. Dot sizes: normalized functional scores for modules on this brain function. Dot colors: functional scores derived from soma distribution of modules (blue); functional scores derived from PBs distribution of modules (red).

### Bouton-net matches arbor-net in cortical, thalamic, and striatal modularity

The arbor-net exhibits well-organized modularity in its eight clusters, A1 through A8. First, the dense module connectivity within these clusters indicates an elevated degree of inter-connectivity among specific neuron groups (**Figure 1B**). Second, these clusters show distinct preferences for cell body locations (**Figure 2A**). Indeed, most cells in clusters A1 and A7 originate from the thalamus (VPM), while the majority in A4, A6 and A8 are from the striatum (CP). Third, the axonal arborization patterns across all clusters are homogeneous as seen in the brain maps (**Figure 2A**).

The bouton-net was constructed by utilizing the segmented PBs from individual neurons (**Methods**). It was not surprising that there would be some differences between bouton-net and arbor-net, but overall, it showed both networks have clear modularity (**Figure 2B**).

Indeed, all three of the above characteristics of the modularity of bouton-net can be clearly seen by illustrating both the incoming upstream sources and outgoing downstream targets of specific brain regions. We delineated the sorted connections of cortical, striatal, and thalamic regions in modules B1 to B12 (**Figure 2C-E**; **Supplementary Figure 4**). In the example of the VISp region (**Figure 2C**), modules B6, and B8 emerge as clear hubs, highlighting a broad range of modules coding the whole-brain connectivity. Cortical neurons in VISp (**Figure 2C**) receive 43.33% of their incoming connections from VISp, a similar amount from other LGd neurons, and a smaller number of upstream connections from other cortical areas. We observed that VISp neurons project more than 60% of their outgoing connections to neurons in the same area, and other visual areas including VISal, VISrl, and VISI. Our data confirms previous reports of LGd-VISp circuits and other findings based on viral tracing for cortical projection mappings, e.g. VISp neurons receiving projections from LGd to mediate visual perception (Bienkowski, et al, 2019; Cruz-Martín, et al, 2014; Douglas, et al, 1989; Douglas, et al, 2004; Jaepel, et al, 2017; Oh, et al, 2014; Sanderson, et al, 1991; Cossell, et al, 2015; Muir, et al, 2017; Harris, et al, 2019).

Neurons in non-cortical regions, such as the striatum and thalamus, also exhibit well-established modularity in bouton-net. For instance, excluding the connections among CP neurons themselves, CP neurons receive most of their incoming connections from cortical neurons such as primary motor area (MOp) but output a substantial portion of their outgoing connections down there (**Figure 2D**). Our data also show that CP neurons project to SNr (substantia nigra, reticular part) or GPe (globus pallidus, external segment), consistent with previous findings of two subclasses of medium spiny neurons (Gerfen, et al, 2011): dopamine receptor D1 (Drd1) neurons projecting to SNr (direct pathway) and dopamine receptor D2 (Drd2) neurons projecting to GPe (indirect pathway).

In contrast, neurons in the thalamic region VPM both receive and output a broader range of connections to cortical regions than CP neurons (**Figure 2E**). Experiments of anterograde and retrograde labeling have demonstrated the reciprocal connections between VPM and widespread cortical regions (Bernardo, et al, 1987; Bourassa, et al, 1995; Graziano, et al, 2008; Lu, et al, 1993; Oh, et al, 2014; Wimmer, et al, 2010; Yang, et al, 2013). Recent neuronal reconstructions and axonal analysis of VPM cells (Peng, et al, 2021; Timonidis, et al, 2023; Winnubst, et al, 2019) also characterized the full diversity of its projections. However, different from our approach, little previous work quantified the neuronal connections within these brain regions.

The ubiquitous modularity of cortical, striatal, and thalamic neurons represents distinct connectivity patterns in the bouton-net clusters. Similarly, we also observed clear modularity in the arbor-net clusters, reflected in their neuronal projection patterns (**Figure 2F**). For instance, clusters A6 and A8 mainly project to the MOp, MOs, and SSp-m regions, but A8 also projects to the RSP (Retrosplenial area), which is different from A6. Clusters A1, A7, and A2 primarily target various cortical regions.

Remarkably, we found strong matches between the modules of the bouton-net and arbor-net (**Figure 1G**). A1 corresponds primarily to B12, but also matches B5, B7, and B10. A6 matches to B1, B3, and B4, jointly. A7 corresponds to B9, B6 and B2. A2 matches to B8 primarily. A8 corresponds to B7 and B11 jointly. The smaller modules A3 and A5 do not show strong quantitative matches to bouton-net modules. In contrast, all bouton-net modules exhibit strong correspondences with arbor-net modules. Our results showed that the bouton-net provided a restrictive set of modules of neuron-level connections for the cortical, striatal, and thalamic regions, and these neuron-level connections among brain regions would not be random.

### Predicted boutons match potential arbor contacts statistically, spatially, and anatomically

Like the brain-wide distribution of PBs (**Figure 1C**), our data indicate that cells originating from the same brain regions tend to have their PACs concentrated in several specific target brain regions, based on extracting 427,206 PACs from F1877 (**Figure 3A**). For example, cortical neurons have abundant PACs in the striatum and thalamus (**Figure 3D**). However, neurons from regions like MOp and MOs also possess PACs in cortical areas. Cells from VPM and VPL exhibit a wide distribution of PACs across various brain regions. This brain-wide distribution of PACs further supports the modularity of the arbor-net observed in above analysis.

We discovered that the anatomical distribution of PBs correlates with that of PACs, even though they were generated completely independently, following different principles. A clear positive correlation (Pearson’s *r* = 0.62, *p* = 1.79e-102) was observed between them across cortical, thalamic, and striatal regions (**Figure 3B**), where we had both PBs and PACs. Additionally, for cells from the same soma-regions, we aggregated and compared their respective counts in the corresponding target brain regions. These two series of counts follow the same trend (**Figure 3B**). This strong correlation found for anatomically corresponding regions suggests that the whole-brain scale estimation of PACs and PBs likely provides valuable information about neuronal connections, more than just random guesses about the locations of potential synaptic sites.

We further investigated whether PBs would colocalize with PACs spatially. It turns out that 99.45% of PBs are within a maximum 3-D distance of 1μm from PACs, contrasting with an even distribution one would expect from random guesses of potential neuron contacts’ locations (**Figure 3C**). As a sanity check, only about 2.74% of PBs are within the 1μm neighborhood of branching points of neurons, which represent another type of morphologically key locations (**Figure 3C**).

We hypothesized that the spatial colocalization and statistical correlation of PBs and PACs would result in co-occurring projection patterns in individual neurons. To explore this, we generated brain-region-sorted whole-brain projection matrices for both PACs (**Figure 3D**) and PBs (**Figure 3E**). We observed that neurons in MOp2/3, MOp5, and MOs2/3 form a closely connected network, in terms of both PACs and PBs. SSp and SSs neurons exhibit more concentrated self-connections, as indicated by the correlated patterns of PACs and PBs. Notably, SSs6a neurons stand out as outliers, differing from other SSs neurons; indeed, they have PAC- and PB-projections to the MOp and MOs areas like those of CLA neurons. CP neurons display unique projection patterns, primarily connecting to themselves. In contrast, VPL and VPM neurons have broad projection patterns extending to many different brain regions. Overall, we found that PACs and PBs share a high degree of projection concentration throughout the entire mouse brain. The remaining difference between PBs and PACs was likely due to the limited number of dendrites used in obtaining PACs, resulting in an order of magnitude fewer PACs compared to PBs (compare **Figure 3A** with **Figure 1C**). To address this, we utilized dendrites from the D18370 dataset (**Supplementary Figure 5**) in conjunction with axons from the F1877 dataset, generating 4,121,201 PACs (see **Supplementary Figure 6A-B**). With this expanded dataset, we found that the anatomical correlation between PBs and PACs improved from 0.62 (**Figure 3B**) to 0.76 (**Supplementary Figure 6C**).

Overall, we found statistical, spatial, and anatomical correlations between PBs and PACs. Such co-occurring patterns are far from random. They offer a high-resolution overview of the neuronal connection network in mouse brains, which had not been achieved through millimeter-scale whole-brain network analysis or nanometer-scale imaging of synapses at the whole-brain scale.

### Predicted neuron connection sites concentrate on functional modules

To complement the brain-wide distribution of PBs (**Figure 1C**) and the map of projection regions (**Figure 3E**), we also produced a geodesic distance map of concentrated PBs (**Figure 3F**). Our data show that most cortical neurons have a high density of PBs within 2mm of their somas. An exception is the cortical CLA neurons, which peak in layer 1 of the ventral part of the Retrosplenial cortex (RSPv1), involved in various cognitive processes such as navigation, memory, and spatial orientation (Vélez-Fort, et al, 2018). Our data seem consistent with previous observations regarding the input-output organization of the mouse claustrum (Zingg, et al, 2018). Unlike cortical neurons, CP neurons exhibit the greatest density of PBs within the CP region, but thalamic neurons evidently have a majority of their PBs in association with various cortical neurons (**Figure 3F**). For instance, neurons in the dorsal lateral geniculate nucleus (LGd) have a peak of 26.62% of PBs found in layers 2 and 3 of the primary visual cortex (VISp2/3). This finding aligns with the vital role of LGd in visual information processing, where the LGd receives input from the retina and sends projections to the primary visual cortex (Bienkowski, et al, 2019). VISp2/3 contains pyramidal neurons essential for intracortical and cortico-cortical communication, involved in tasks such as pattern recognition, depth perception, and motion detection (Ji, et al, 2015).

To further explore the regional projection and enrichment of PBs in terms of various functions, we produced a PB-function map to better understand the relationship between brain region functions and the bouton-net modules (**Figure 3G**). We discovered that the relatively dense connections within each module correspond to anatomical areas responsible for specific brain functions. We evaluated the involvement of all modules in specific brain functions based on the soma distribution of these modules as well as their PB target regions (**Figure 3G**; **Methods**). Our data show that each bouton-net module contributes to at least one type of brain function (**Figure 3G**). For example, modules B6, B8, and B11 are associated with vision and predominantly consist of cells originating from VISp and LGd.

This PB-function map also highlights several other interesting observations (**Figure 3G**). Our data show that functions related to motion and feeling are ubiquitous across brain-wide neuronal connections. In contrast, sensory functions such as smell, taste, vision, and hearing correspond to specific neuronal connections that are enriched in different brain regions and connection modules. Indeed, the bouton-net modules related to sensory functions partially complement each other, forming a conjugate code. Additionally, our data indicate that the connections related to memory and learning are highly correlated, yet they exhibit a dramatically lower correlation with the bouton-net modules associated with emotion, attention, and cognition. Interestingly, the module related to sleep, labeled B11, is involved in almost all functions, except for learning and non-vision sensory functions, including smell, taste, and hearing.

### Single-cell neuronal connections correlate with mesoscale neuronal connectomes and relative density of brain-scale synapse distribution

Previously, a mesoscale connectome, conveniently referred to as ‘Allen-net’, was reported based on visualizing 3-D registered and multiplexed viral tracing at the whole-brain scale (Oh, et al, 2014). The injection sites were much larger than single cells, resulting in neuron populations, often containing hundreds or thousands of neurons, being mapped in this connectome. Allen-net has been a valuable reference for interconnected brain regions due to the common use of anterograde (e.g., adeno-associated virus and herpes simplex virus) and retrograde (e.g., modified rabies virus and pseudorabies virus) tracing viruses injected into different brain regions for neural circuits. In addition, a more recent mesoscale connectome, referred to as ‘BRICseq-net’, of brain regions was derived from a single cell barcoding and sequencing technique (Huang, et al, 2020). Thus, a direct comparison of our single-neuron connectomes against Allen-net and BRICseq-net is useful to understand whether the single-cell connections will add useful knowledge to connectome studies.

As in their original studies, both the Allen-net and BRICseq-net were represented at the level of brain-regions, for the purpose of a head-to-head, region-wise comparison, we reduced the resolution of our connectomes to either 76 (for Allen-net) or 33 (for BRICseq-net) common brain regions throughout a mouse brain (**Figure 4**). To do so, we multiplied the corresponding connection strength scores in both the arbor-net and the bouton-net, aggregated all scores within the respective brain regions, and combined them into a single meso-scale connectome called ‘SEU-net’ (**Figure 4A**; see **Supplementary Figure 7** for a complete set of 148 brain regions including cortical layers; **Methods**). Both the total numbers of PACs and PBs as used in constructing SEU-net have a strong positive correlation with the connection strength of the Allen-net and BRICseq-net (**Figure 4B**), for the shared brain regions with available data. However, as the connection strength of SEU-net is defined based on the average number of PACs and PBs (**Methods**), there is a weak correlation (*p* = 0.014) between the connection strength of SEU-net and Allen-net, whereas the correlation between the connection strength of SEU-net and BRICseq-net is still strong (*r* = 0.26, *p* = 2.90e-6) (**Figure 4C**). Overall, the connections of SEU-net and previous work as shown in Allen-net and BRICseq-net are mostly consistent.

**Figure 4.**
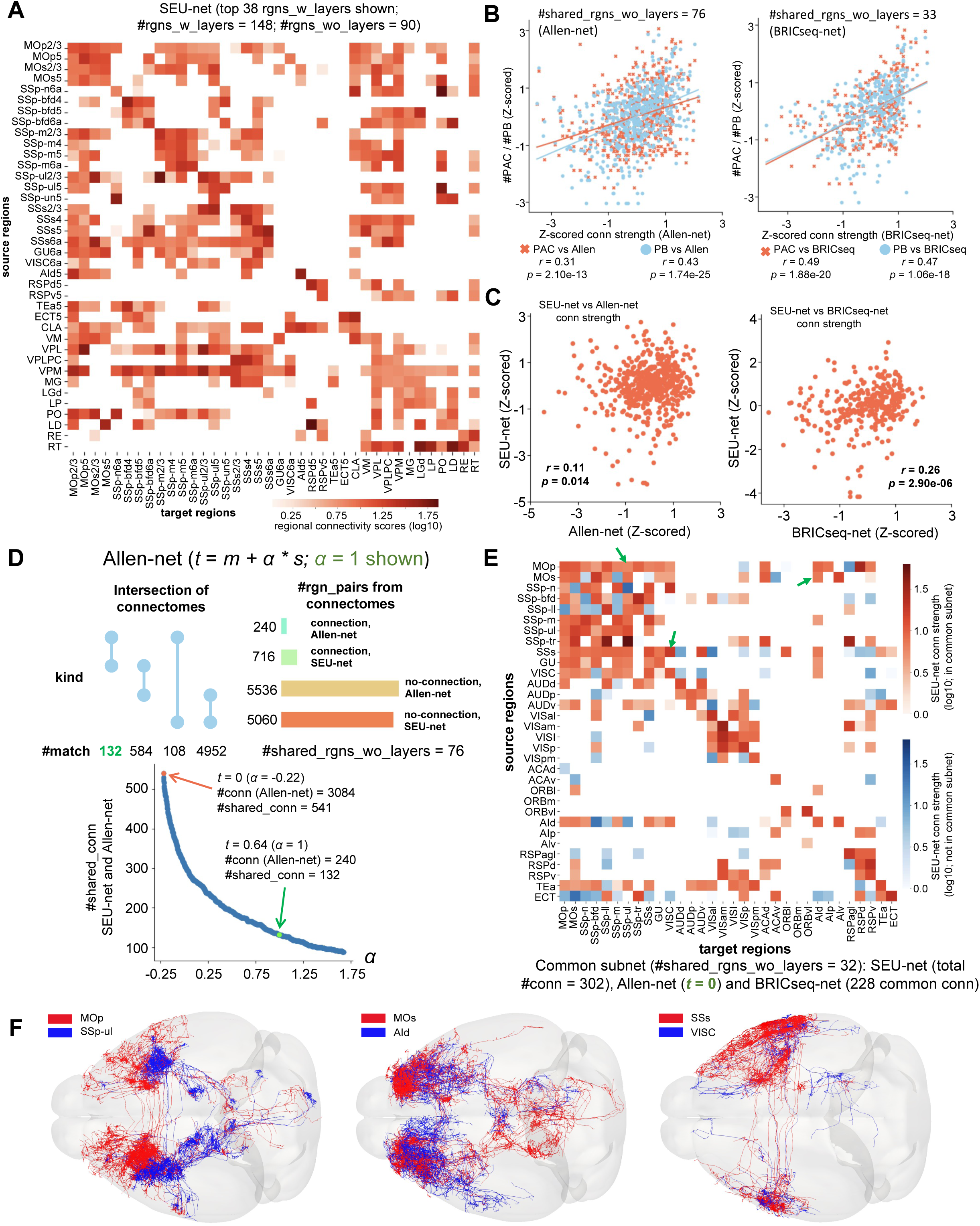
Cross-validation of three meso-scale connectomes. **A.** Regional SEU-net: regional connectivity map (within 38 brain regions with cortical layers). Vertical axis: 38 source brain regions of the highest outgoing connection strength. Horizontal axis: 38 target brain regions same as vertical axis. Color bar: connection strength (log10-transformed) between source and target region. **B.** Correlation between mesoscale connection strength (Allen-net and BRICseq-net) and anatomical distributions of PACs (cross, red) and PBs (dot, blue). Vertical axis: counts of PBs and PACs. Horizontal axis: z-scored connection strength (left: Allen-net; right: BRICseq-net). **C.** Scatter plots of corresponding connection strength between mesoscale connectomes (SEU-net, Allen-net and BRICseq-net). Vertical axis: z-scored connection strength of SEU-net. Horizontal axis (left): z-scored connection strength of Allen-net. Horizontal axis (right): z-scored connection strength of BRICseq-net. **D.** Intersection of SEU-net and Allen-net. Top: counts of connections and non-connections of SEU-net and Allen-net within 76 shared brain regions (left), and their intersections (right). Bars represent the counts of connections and non-connections. Dots and lines indicate the types of intersections. Bottom: dynamic thresholding of the Allen-net to determine connections and non-connections. Vertical axis: counts of shared connections in both connectomes. Horizontal axis: parameter to determine the threshold. **E.** The common subset of three mesoscale connectomes. Color bars indicate the connection strength of SEU-net (red: connections in common subset; blue: connections not in common subset;). Arrows annotate three connection examples. **F.** Examples of neuron connections in SEU-net. Three groups of single-neuron level connections (red: source neurons; blue: target neurons) are displayed in CCFv3. The regional connections are also marked in Fig. 4E with green arrows.

Another way to cross-validate different connectomes is by intersecting their connections and non-connections. In the case of Allen-net, there is a connection strength score between each pair of brain regions. Therefore, we compared the common connections of SEU-net and Allen-net, applying a dynamic threshold to Allen-net (**Figure 4D**). 75.6% of SEU-net connections (541 out of 716) were found in Allen-net when considering all connections in the latter. For the statistically significant connections in Allen-net, whose connection strength is greater than one standard deviation above the respective mean strength value, 55.0% of the predicted connections in SEU-net (132 out of 240) were confirmed by Allen-net. At the same time, 97.9% of non-connections observed in SEU-net (4952 out of 5060) were also confirmed by Allen-net (**Figure 4D**). This ballpark consistency between the two connectomes seems reasonable, given that Allen-net is known to contain false-positive connections but is unlikely to have many false negatives. The remaining differences between Allen-net and SEU-net could be due to different neurite labeling techniques, imaging protocols (e.g., sensitivity), and data processing (e.g., registration methods) (see **Discussion**).

We further cross-validated all three connectomes, i.e. SEU-net, Allen-net, and BRICseq-net, by finding their greatest intersection (**Figure 4E**). To do this, we considered the 32 common brain regions of these three datasets. We found that most SEU-net connections were present in their common subnet of three regional connectomes (**Figure 4E**). For instance, SEU-net’s aggregated predicted connections from MOp neurons to SSp-ul neurons, from MOs neurons to AId neurons, and from SSs neurons to VISC neurons (**Figure 4F**) were all confirmed at the brain region level, as shown in the common subnet of the three connectomes (**Figure 4E**). This result indicates that our approach to assembling a single-neuron connectome is more likely to be useful than a random guess, and single-neuron connections correlate with meso-scale neuronal connectomes.

In further cross-validation, we compared the potential connections predicted by our approach with a recent brain-wide synaptome (Cizeron, et al, 2020). We analyzed the average synapse distribution from 11 mouse brains aged between 60 and 99 days, naming the dataset Grant2020 after the senior author (Cizeron, et al, 2020). We identified 102 common brain regions between our predicted brain (PB) regions and the regions studied in Grant2020. Across these 102 regions, we observed a strong correlation (*r* = 0.49, *p* = 2.33e-7) between PBs and the relative synapse distribution (**Supplementary Figure 8A**). There was also a positive correlation between predicted axonal connections (PACs) and Grant2020 (**Supplementary Figure 8B**). Coupled with the consistent patterns found across three connectomes regarding the incoming and outgoing connections (**Figure 4E**), it appears that the spatial distributions of PACs and PBs (**Figure 3**) capture key aspects of the spatial granularity and organization of potential synaptic connections.

We also performed a survey of existing literatures concerning 156 sampled connections from the common subnet of three regional connectomes (**Figure 4E**). It turns out that 87.18% of the sampled connections (136 out of 156) were previously studied, demonstrating regional connections in terms of anatomical structures or functional activities, complementing the validation of our neuronal connections. The literatures and evidence were documented in **Supplementary Table 2**.

### Single-cell neuronal connections correlate with co-expressed genes, outperforming neuron-population based mesoscale connections

It remains an open question whether neuronal connections are supported by gene co-expression, as the co-expression of genes in different brain regions only indicates a possibility that these regions might be connected by a neuronal pathway (e.g., Hartl, et al, 2021). While one might hypothesize that a greater number of genes shared in co-expression patterns would imply a higher likelihood of some kind of association of brain regions, either through direct or indirect neuronal connections, validating such a hypothesis has been limited by the lack of specifically designed experiments in this field. This challenge is further complicated by the fact that many genes, ubiquitously expressed in the brain, present complex background co-expression patterns, even when there is no direct neuronal connection between brain regions.

In a brain-wide analysis, we combined SEU-net with the genome-wide, *in situ* hybridization-based Allen Brain Atlas (ABA) gene expression database (Lein, et al, 2007) to generate a region-wise gene co-expression map for 19,907 mouse genes (**Figure 5A** showing selected 32 layer-merged regions; **Supplementary Figure 9** showing all 76 brain regions). Upon visual inspection, we observed a clear correlation between these co-expression maps (**Figure 5A**) and the regional connection map, arranged in the same order (**Figure 4E**). To quantify this relationship with examples, it is noteworthy that the relatively strong connection from MOp to SSp-m corresponds to 32.12% of co-expressed genes from the entire mouse genome, with an average gene correlation score of 0.60 (**Figure 5B**; **Methods**).

**Figure 5.**
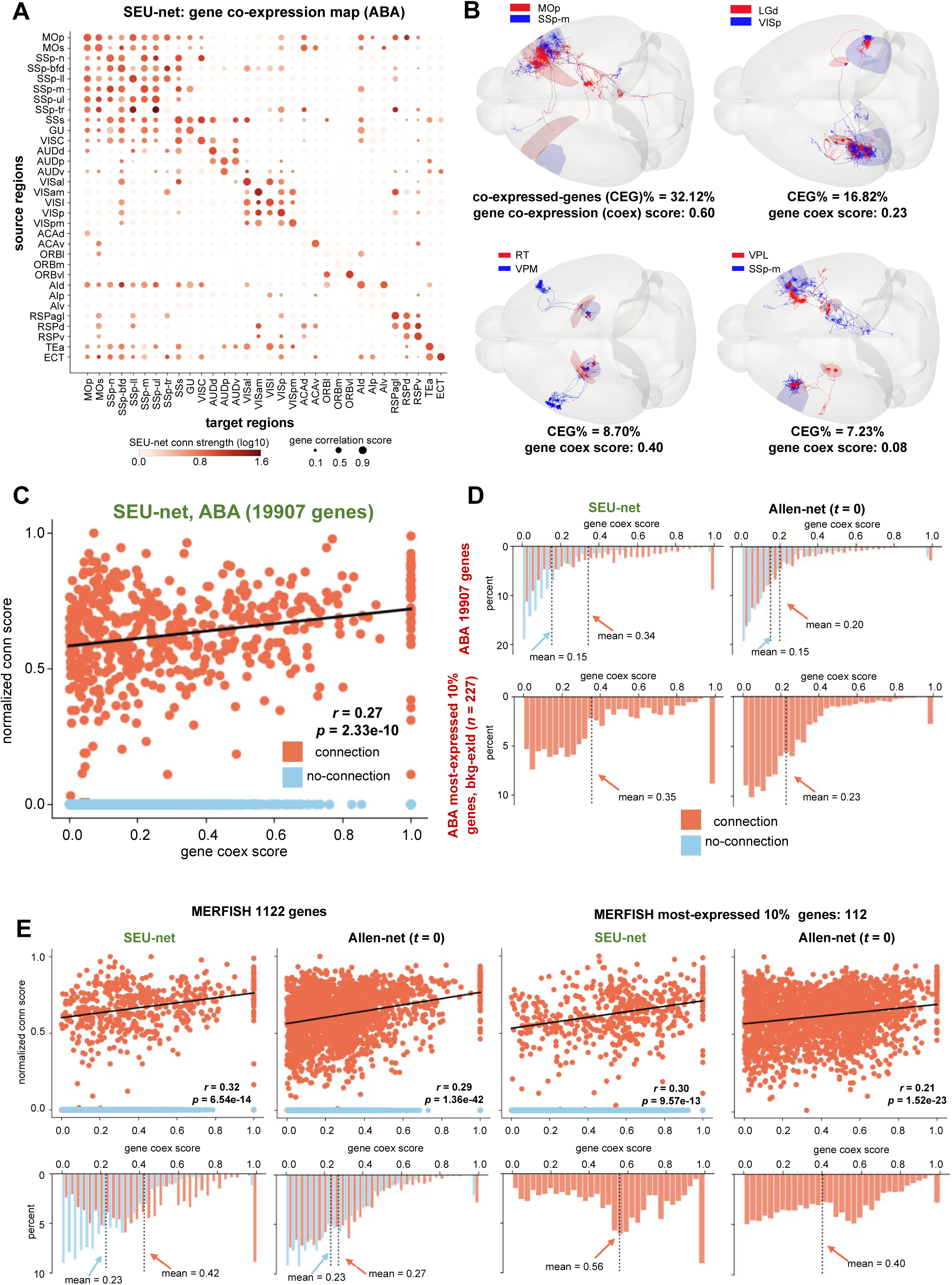
Validation of neuronal connections using genome-wide gene co-expression and single-cell MERFISH. **A.** Whole-brain region-wise gene co-expression map using ABA dataset. Vertical axis: 32 source brain regions. Horizontal axis: 32 target brain regions. Dot colors: regional connectivity of SEU-net. Color bar: regional connection strength from SEU-net. Dot sizes: gene co-expression scores. **B.** Regional connectivity and gene co-expression examples. Red: source neurons; blue: target neurons. **C.** Scatter plot of corresponding regional connection strength and co-expression scores within all shared 76 brain regions (between SEU-net and ABA dataset; red: connected brain regions; blue: non-connected brain regions). **D.** Distributions of gene co-expression scores for connections and non-connections from both SEU-net and Allen-net (t=0). Horizontal axes: gene co-expression scores. Vertical axes: percentages of connections (red) and non-connections (blue) in each bin. Top row: using all 19907 mouse genes in ABA dataset. Bottom row: using 227 ‘non-background’ genes. Left column: using connectivity from SEU-net. Right column: using connectivity from Allen-net. Red arrows indicate the average gene co-expression scores for connections, and blue arrows indicate the average gene co-expression scores for non-connections. **E.** Scatter plots and distributions of corresponding regional connectivity and gene co-expression using MERFISH dataset. Left two columns: using all 1122 genes. Right two columns: using the 10% most expressed genes. Each column uses regional connectivity from SEU-net and Allen-net respectively. Arrows indicate the average gene co-expression scores in the same way in **D**.

Quantitatively, for the neuronal connections analyzed using SEU-net, we found a clear correlation (*r* = 0.27, *p* = 2.33e-10) between the strength of neuronal connections and the level of gene co-expression (**Figure 5C**). Additionally, we profiled the histogram of gene co-expression scores associated with specific regional connections (**Figure 5D**). We discovered that the genome-wide gene co-expression of connected brain regions exhibited a longer tail compared to that of non-connected regions, as indicated by both SEU-net and Allen-net connectomes. However, the gene co-expression for non-connected regions displayed similar histograms. This observation is consistent with the hypothesis that ubiquitously expressed genes contribute to a background co-expression pattern. Therefore, we also analyzed the most highly expressed ‘non-background’ genes, after excluding all ubiquitously co-expressed ‘background genes’ (**Figure 5D** - bottom row). In both analyses, connections identified by SEU-net consistently showed higher average gene co-expression scores than those identified by Allen-net. For instance, when considering the top 10% most highly expressed genes, after removing the ubiquitous background, the mean gene co-expression score for SEU-net connections is 0.35, which is 52% higher than that for Allen-net connections (0.23) (**Figure 5D** - bottom row). Our findings suggest that the single-neuron reconstructions used to construct SEU-net are more strongly correlated with gene co-expression than the neuron population-based Allen-net.

We also cross-validated SEU-net with an alternative single-cell gene expression dataset, which profiled 1,122 expressed genes using a Multiplexed Error-Robust Fluorescence in situ Hybridization (MERFISH) technique (Zhang, et al, 2023). We generated a region-wise gene co-expression map based on the MERFISH data (**Supplementary Figure 10**), displaying an apparent correlation with our genome-wide co-expression map (**Supplementary Figure 9**). More interestingly, although the background ubiquitously co-expressed gene histograms drawn from the combined analysis of SEU-net, Allen-net, and the MERFISH data (**Figure 5E** – left two columns) still exhibit a monotonously decreasing profile, similar to those observed in the respective ABA analysis (**Figure 5D**), the histograms for strongly co-expressed genes in SEU-net show a bell-like profile (mean = 0.42), differing from that of the Allen-net (mean = 0.27) (**Figure 5E** – left two columns). With the top 112 most expressed genes, SEU-net connections correspond to a mean co-expression score of 0.56, which is 40% higher than that of the Allen-net (mean co-expression score = 0.40) (**Figure 5E** – right two columns). Additionally, we note a consistent, positive correlation between gene co-expression scores and brain region connections, whether in SEU-net or Allen-net. In other words, we demonstrate the quantification of the approximately linear correlation between neuronal connections and gene co-expression, whether the gene expression data is genome-wide or single-cell-oriented. Furthermore, the single-neuron connections as depicted in SEU-net correlate more distinctly with gene co-expression, using different methods, than the neuron-population-based connections revealed in the Allen-net.

### Single neuron connectomes feature consistent subnetwork motifs and other network properties

After analyzing the global connections in both the arbor-net and bouton-net, which are likely real, we further examined whether the combination of subsets of these connections would also make sense. To do this, we studied a series of network properties of the single neuron connectomes and their derivatives, specifically focusing on the local network structures and characteristics of the arbor-net, bouton-net, and SEU-net.

The detection of subnetwork patterns, i.e., motifs, in complex networks is a widely adopted method for studying the intricate relationships among network nodes and their connections (edges). There are various types of network motifs, such as dyads, triads, and tetrads, involving 2, 3, and 4 network nodes, respectively (Milo, et al, 2002; Alon, 2007; Felmlee, et al, 2021). In this study, we focused on triads that cannot be directly deciphered using bivariate analyses of pairs of nodes. We counted the occurrences of 13 triad motifs (M1 ~ M13) (**Figure 6A** – top). Since each triad involves a different number of directed edges between nodes (for example, M1 involves two directed edges, and M8 involves four directed edges (**Figure 6A** – middle)), it is natural that triads with more directed edges would have fewer occurrences. Therefore, we normalized the observed occurrences using the respective expected values in random networks (**Figure 6A** – bottom) and discovered outstanding triads representing fundamental local subnetwork patterns involving three nodes in arbor-net, bouton-net, and SEU-net connectomes (**Figure 6A** – bottom, boxed triads). The observation that triads featuring a greater number of directed edges are more abundant underscores the presence of intricate subnetwork patterns within these connectomes. This supports our hypothesis that an analysis of these connectomes should extend beyond global bivariate analyses to include comparisons within the motif space, thereby providing more understanding of their structural complexities. In particular, we found a few motifs, e.g. M9 triads, were overabundant in both arbor-net and bouton-net. Thus, we profiled them more closely as follows.

**Figure 6.**
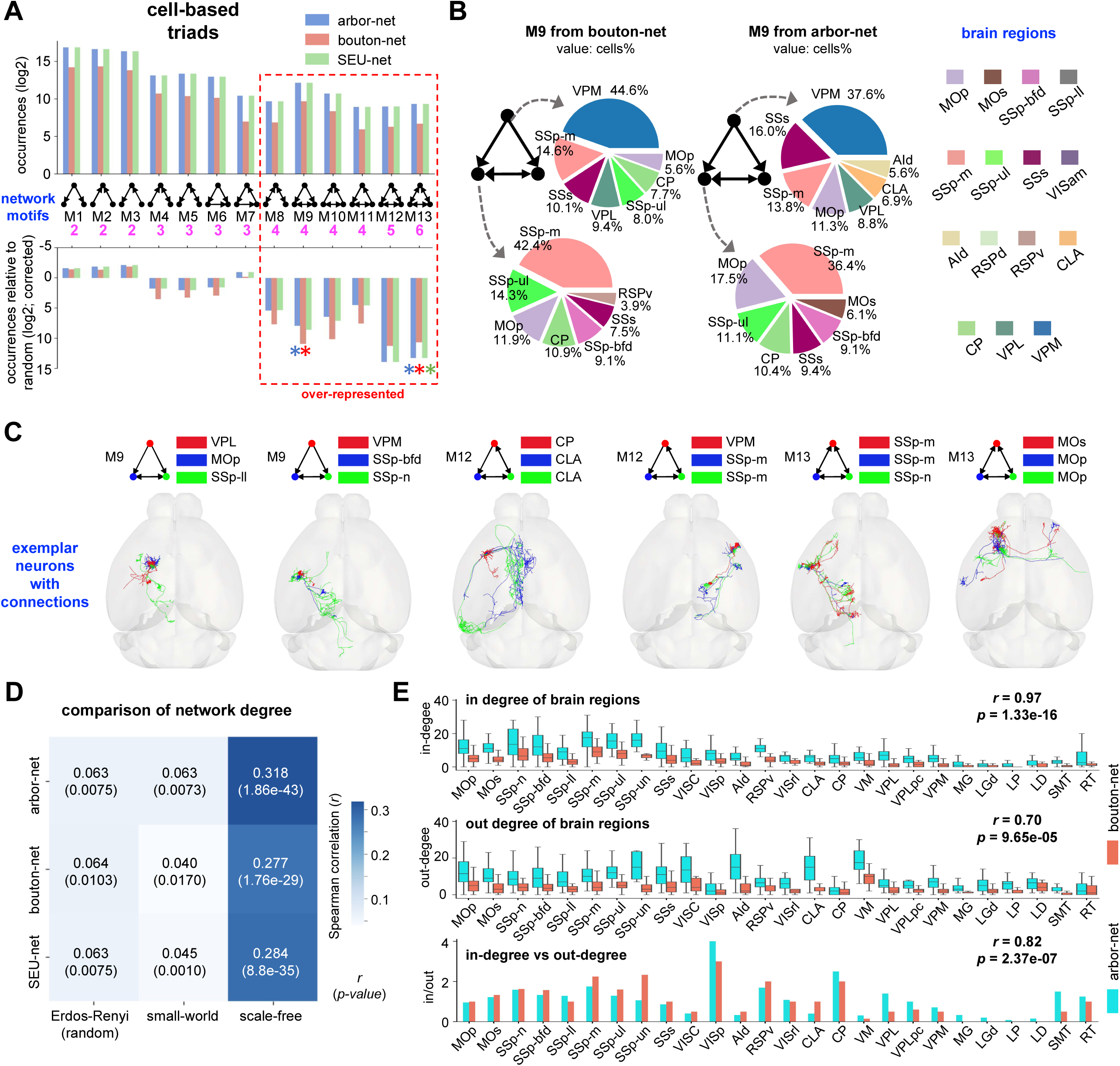
Network analysis of the single neuronal connectomes indicates consistent subnetwork (motif) patterns and other network properties. **A.** Network motif distributions of connectomes and comparison with the random Erdos-Renyi (ER) networks. Top: occurrences of motifs from arbor-net (blue), bouton-net (red), and SEU-net (green). Colors indicate the respective connectomes. Middle: structures of network motifs and their counts of edges. Bottom: corrected motif occurrences relative to ER network with same nodes and same edges. Box and stars highlight the over-expressed motif patterns of connectomes. **B.** Statistics of neuronal sources in motifs across brain regions. Pie-charts represent nodes (arbor-net and bouton-net: neurons) of motifs, and arrows represent their edges (predicted connections). Texts around the pie-chart indicate the percentages from corresponding source brain regions. **C.** Exemplar neurons and their connections in triads M9, M12, and M13. Top: motif structures formed by nodes and edges, and node colors corresponding to their soma locations. Bottom: neuron reconstructions in CCFv3. **D.** Comparison of degree distribution between connectomes and random networks. Values in matrix: Spearman correlation coefficients and p-values between degree distributions, also annotated by texts in the heatmap. **E.** Distributions of in-degree and out-degree in major brain regions from arbor-net (blue) and bouton-net (red). Top: box plots of in-degree distributions. Middle: box plots of out-degree distributions. Bottom: box plots of in-degree versus out-degree. Horizontal axis: 26 brain regions, each with >10 cells. Vertical axis: degree values (top two), and ratios of in-degree to out-degree (bottom). Texts (top right in each bar plot) indicate the Pearson correlation coefficients and p-values between arbor-net and bouton-net.

Our analysis revealed that the two M9 motifs from arbor-net and bouton-net, despite being derived independently, exhibited remarkably similar profiles regarding the source and target brain regions of the underlying neurons (**Figure 6B**). More precisely, the nodes at the apex of the arbor-net M9 and the bouton-net M9 both were predominantly composed of cells from the VPM, SSp-m, SSs, VPL, and MOp regions. The nodes at the base of the two M9 triads both predominantly comprised cells from the SSp-m, SSp-ul, MOp, CP, SSp-bfd, and SSs regions. The proportional distribution of these cells within the two sets of triads’ profiles also had a clear resemblance to each other. This pronounced correlation between subnetwork patterns that were derived independently underscores that our single neuronal connectomes are characterized by non-random network connections and structures, which can be consistently identified using varied methodologies. We also observed similar situation for other motifs, such as M8 and M10 ~ M13 motifs (**Supplementary Figure 11**).

In addition to the motif analysis, we also examined other network properties. A previous study on Allen-net (Oh, et al, 2014) showed that the degree distribution of the mouse brain network best approximates that of a scale-free network. Our tests on both single neuronal connectomes, i.e. arbor-net and bouton-net, as well as the regional connectome, i.e. SEU-net, yielded similar conclusions (**Figure 6D**). We also found that arbor-net and bouton-net shared substantially correlated in-degree and out-degree profiles, as well as their ratios (**Figure 6E**). All these pieces of evidence support the notion that our single neuronal connectomes carry brain-wide global properties of single neuronal connections that are non-random and can be cross-validated using state-of-the-art, multi-modality datasets from independent sources.

## Discussion

Understanding single-cell level connectivity across the entire brain is crucial for comprehending its structural wiring and functional activities. Many studies on the long projection morphology reconstruction of single neurons have been published in recent years (Winnubst, et al, 2019; Peng, et al, 2021; Gao, et al, 2022; Liu, Yun, et al, 2023; Liu, Jiang, et al, 2023; Qiu, et al, 2024), enabling investigations into regionally connectivity at the single-cell level through axonal projections. However, these works have scarcely delved into the specifics of how two neurons are probabilistically connected to each other, due to a variety of challenges, e.g. some of the published datasets do not include complete dendrites and axons at the same time. Our study makes an initial attempt to provide a practical and reliable framework for mapping neuronal connections between individual neurons, using one of the largest available neuron reconstruction datasets (Liu, Jiang, et al, 2023).

In particular, we have developed an extensible framework for constructing connectome models of single neurons reconstructed at the whole-brain scale. We have also provided a comprehensive set of cross-validation studies to justify the statistical significance of the connectomes and findings obtained using our approach. Essentially, our paradigm builds detailed connectome models of individual neurons by mining 3-D reconstructed and registered neuron morphologies in a standardized atlas space. Our approach first quantifies the spatial adjacency of paired axonal-dendritic arbors. Complementing this, we have introduced an independent method that utilizes information from individual neurons — not just their pairs — by quantifying the predicted axonal varicosities (conveniently called “boutons”), based on the axonal morphologies reconstructed at the whole-brain scale. We have cross-validated these two methods in multiple ways, particularly through the creation of brain network models, namely an arbor-net and a bouton-net, and their respective modules. We observe that these modules exhibit strong granularity, characterized by sparse cross-module connections but relatively abundant intra-module connections throughout cortical, striatal, and thalamic regions.

Remarkably, these two independently developed brain network models demonstrate strong statistical consistency, which is highly unlikely to arise from mere conjectures of potential neuronal connections. We further validate our approach using independently established other multi-level, multi-modality datasets, including synapse distribution, gene expression, and mesoscale connectomes, as well as very intensive literature search. We have found that most connections consolidated based our detailed connectome models can match established knowledge in literature, but at the same time we also discover many previously unknown connections that deserve future examination. Therefore, our paradigm establishes a practical and extensible framework for building single-neuron connectomes using increasingly available 3-D morphological reconstructions of neurons, without the need for — but apparently can be enhanced by — additional nanometer-scale imaging of synapses, which is not yet feasible for whole brains. We believe the generated connectomes capture the indispensable properties of brain networks, and our approach provides an instrumental framework as more individual neurons’ morphologies become accessible.

Traditionally, the term ‘Peters’ rule’ (Braitenberg and Schüz, 2013) was used to describe the method that estimates the strength of potential synaptic connections based on the spatial adjacency or overlap of axonal and dendritic arbors (Rees, et al, 2017). This intuitive approach to approximating neuronal connections is appealing, as neurons whose arbors do not overlap are unlikely to connect directly. However, it is also true that spatial adjacency does not guarantee a connection, as it could be due to passing neurites without actual synaptic connections, among other complications. Nevertheless, it is fair to say that spatial adjacency does increase the likelihood of connectivity. Such probabilistic, predicted connectivity is very useful for understanding the overall brain networks. Additionally, systematic profiling of individual neurons’ probabilistic connectivity has recently been shown to be fundamentally crucial and powerful for classifying neurons into connectivity subtypes that could not be discovered using morphology alone (Liu, Yun, et al, 2023). Despite these successes, a key unanswered question remains: how can one ascertain whether spatially adjacent or overlapping neurons are likely to form real connections? Our approach in this paper offers a direct and original answer to this key question. Essentially, our argument is that individual neurons’ axonal varicosities can complement the spatial adjacency of pairs of neurons with overlapping axonal-dendritic regions, producing a more realistic estimation of connection probability. This basically leads to an extensible, new way to predict neuronal connectivity that can be cross-validated using independently produced evidence from different data modalities, as elaborated in this paper. In other words, one of our key points is that Peters’ rule alone is not sufficient; it must be conjugated with other evidence and can be conveniently enhanced by using statistics on putative axonal boutons, as we have shown.

Although we have cross-validated our connectomes using an independently produced synaptome (Cizeron, et al, 2020), we acknowledge that nanometer-resolution imaging (e.g. EM (Yin, et al, 2020; Turner, et al, 2022; Shapson-Coe, et al, 2024)) of synaptic connections in specific brain regions would provide additional verification of the predicted neuronal networks in our current connectomes. It should be noted that while reconstructing a nanometer-scale connectome of mammalian brains using electron microscopy from scratch presents multiple challenges in the sample preparation and time cost, developing a verification process as described here would be a practical approach to achieving a similar objective.

## Acknowledgment

This project was mainly supported by a Southeast University initiative for neuroscience awarded to HP. We thank Yufeng Liu, Zhixi Yun, Shengdian Jiang, Penghao Qian, Yimin Wang, Zuohan Zhao for assistance in providing supporting data and discussion with a number of other colleagues.

## Author Contribution

HP conceptualized and designed this study, and managed the entire project. FX and LL generated the analysis results along with biological interpretation under the detailed instruction of HP. HP wrote the manuscript with input from all coauthors.

## Data and Materials Availability

The data of 1,877 full neuron reconstructions along with predicted axonal boutons, and dendritic reconstructions of 18,370 neurons were archived at https://zenodo.org/doi/10.5281/zenodo.10677899. The generated connectomes are also archived on the repository. Supporting data tables are also provided as **Supplementary Tables.**

## Methods

### Sources of experimental data

We built connectomes based on 1,877 fully reconstructed single neuron morphologies, referred to as “F1877”, derived from our recent work (Liu, Jiang, et al, 2023 - data version 7-24-2023). The dendritic morphologies of 18,370 neurons (referred to as ‘D18370’) and 2.57 million Predicted Boutons (PBs) were also accessed from the same work. In specific, to generate PBs for each neuron, axonal skeletons were first segmented into 20 µm length fragments, where intensity and radius profiles were calculated. Overlapped peaks in both intensity and radius profiles were identified as initial candidate boutons, and false positive results were filtered out with heuristic criteria for bouton size and image intensity (Liu, Jiang, et al, 2023). All neuron morphologies and PBs were spatially registered to the Allen Mouse Brain Common Coordinate Framework v3 (Wang, et al, 2020) using the mBrainAligner tool (Qu, et al, 2022). Subsequently, we resampled these registered neuron morphologies at 1 µm intervals, using the “resample_swc” function in Vaa3D (version 3.601). Summaries of these 20,247 neurons were provided in **Supplementary Table 3**.

### Quantification of neuronal connection score

To quantify the connection scores between neurons, we computed the closest Euclidean distance from axon arbors to dendritic arbors. Considering the typical length of dendritic spines and the size of axonal boutons (Iascone, et al, 2020), we designated the axonal locations within 5μm to the dendrite as PACs. For an axon-dendrite pair with *N* PACs, we defined the connection score using the following equation:

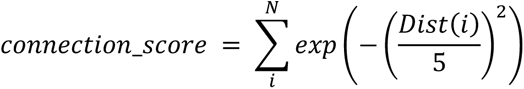

where *Dist*(*i*) represents the shortest Euclidean distance from the *i^th^* PAC to the dendritic arbors. We estimated the connection scores for the F1877 cells, and constructed the arbor-level connectome (arbor-net). Following the same definition of connection scores, we also generated the bouton-level connectome (bouton-net) using the spatial closeness between PBs and target dendritic morphologies, which constrained the potential synaptic contacts within our previously predicted axonal boutons.

### Graph partition algorithm

To further investigate these neuronal connections, we applied Spectral Clustering (with parameters: affinity=“precomputed”, assign_labels=“discretize”, random_state=200) from the scikit-learn Python package (version 1.2.2) to cluster neurons based on their log10-transformed connection scores for each connectome respectively. The optimal number of clusters was determined by the modularity score (using the NetworkX Python package, version 3.0.0) automatically. To ensure the quality of clustering results, we additionally filtered out clusters containing fewer than 30 cells.

### Correlation analysis between bouton modules and brain function

First, we engaged experts to annotate the functionality of brain regions involved in this study (See **Supplementary Table 4** for the list of functionality of brain regions). Subsequently, we weighted these brain functions associated with bouton-net modules based on the proportion of neuronal somas from each module present in these regions, assigning functional scores to each module. Assuming somas from module *C* are distributed across *N* brain regions, the functional score of module *C* related to a specific function *F*, is computed using following equation:

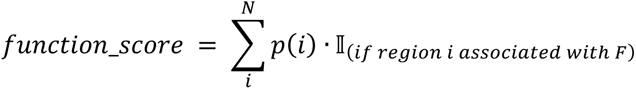

where *p*(*i*) is the proportion of neuronal somas from module *C* in the *i^th^* brain region, and the 𝕀 is indicator function, yielding 1 when the input within the parentheses is true and 0 otherwise. To reveal the clear pattern between these functions and bouton-net modules, functional scores less than 0.05 were set to 0.

### Inter-regional connectivity and comparison with previous brain connectivity atlas

We produced a whole-brain inter-region connectivity map, called SEU-net, and compared it with two previous connectomes, Allen-net (Oh, et al, 2014) and BRICseq-net (Huang, et al, 2020). To estimate regional connection strength of SEU-net, we first multiplied the corresponding connection strength of each neuronal connection from the arbor-net and bouton-net, generating their merged connectome. Subsequently, we averaged the connection strength values of all neuronal connections between source region and target region to represent connection strength of the two regions.

The Allen-net provided quantitative connection strength values involving 213 anatomical regions on both ipsilateral and contralateral hemispheres (Table S3 in Oh, et al, 2014). We multiplied the connection strength of two hemispheres and selected 76 common brain regions in both connectomes for comparison. To determine connections and non-connections, we employed a dynamic threshold *t* in the Allen-net. The threshold was computed as the mean strength value (*s*) plus the standard deviation (*m*) multiplied by a hyper-parameter α.

The BRICseq-net provided brain region-to-brain region connection strength in mouse neocortex from six BRICseq brains (Table S4 in Huang, et al, 2020). The inferred connection strength of each brain was provided in both ipsilateral and contralateral hemispheres. We averaged the connection strength of six BRICseq brains, and then multiplied the connection strengths of two hemispheres. For comparison, we selected 33 common brain regions shared by both our SEU-net and BRICseq-net.

The comparison among the PACs, PBs, and related connectomes was assessed using the Pearson correlation coefficient (scipy; version 1.21.0).

### Whole-brain connectivity and gene correlation

We investigated the relationship between whole-brain connectivity and gene co-expression. The regional connection strength scores of SEU-net (with 76 layer-merged brain regions) were normalized to a range from 0 to 1 by MinMax scaling (scikit-learn Python package; version 1.2.2). Gene co-expression indicates shared genes expressed across brain regions. The co-expression score was defined as the absolute Pearson correlation coefficient (scipy; version 1.21.0) of gene expression strength between two brain regions. The ratio of co-expressed genes was defined as the proportion of co-expressed genes between two brain regions relative to the total expressed genes in both brain regions.

We first acquired two spatial gene expression datasets: the Allen Mouse Brain Atlas (ABA atlas; Lein, et al, 2007), and a recently released spatial transcriptomic cell atlas of the whole mouse brain (MERFISH atlas; Zhang, et al, 2023). The strength of gene expression in brain regions can be derived from the two datasets, and used for the estimation of gene correlations between brain regions. All genes used in the study were documented in **Supplementary Table 5**.

The ABA atlas offers a genome-scale collection of cellular-resolution gene expression profiles using *in situ* hybridization technique. It encompasses the expression patterns of 19,903 genes across the entire brain, derived from 21,553 experiments of sagittal perspective. Each experiment provided “expression energy” of each brain region to represent the regional gene expression strength. To determine the specific gene expression strength of brain regions, we normalized the “expression energy” values of all regions as percentages for each experiment, and then averaged these percentages across experiments for the same genes and brain regions. To exclude the background genes, we first removed all co-expressed genes in non-connected regions, and then selected the top 10% most expressed genes according to their total gene expression strength across all brain regions.

The MERFISH atlas provides a spatial transcriptomic dataset of more than 1,100 genes and approximately 10 million cells across entire mouse brains. The processed MERFISH dataset was organized and stored into four subsets based on their source animals. To avoid potential confounding factors from additional data integration procedures, we selected one subset (log2-transformed count matrix; https://alleninstitute.github.io/abc_atlas_access/descriptions/Zhuang-ABCA-3.html) of 1,122 genes and 606,170 cells (excluding cells not from the 76 common brain regions). Subsequently, the gene expression levels of brain regions were quantified by averaging the single-cell gene counts within the same region, and the top 10% genes of the MERFISH atlas were determined by ordering the total counts of each region.

### Network analysis

In network analysis, neurons are considered as nodes, and their connections represent edges. It enabled us to examine network characteristics such as motifs, and degree distributions of arbor-net, bouton-net, and SEU-net. Here the SEU-net referred to the cell-to-cell merged connectome of arbor-net and bouton-net.

Motifs were fundamental subnetwork patterns in network analysis, and we explored the motifs of 13 triads (M1-M13) within our connectomes. The occurrences of these triads were calculated using the “get_subisomorphisms_lad” function in the iGraph Python package (version 0.11.3).

To identify the most significant motifs in the connectomes, we compared the occurrences of 13 motifs in respective connectomes to their occurrences in random networks accordingly. We independently generated 100 random Erdos-Renyi networks (ER network; Erdos, et al, 1984), and calculated the average occurrence of each motif across these 100 random networks. The normalized motif importance was defined as the result of dividing motif occurrence in each connectome by its average occurrence in random networks and subtracting its number of edges for normalization.

We defined in-degree of a neuron as the number of connections received by the neuron as a target cell, and out-degree as the number of connections originating from the neuron as a source cell. While we constructed our connectomes into “Graph” objects using the NetworkX Python package (version 3.0.0), the in-degree and out-degree of each node were directly computed as inner attributes of the objects.

We conducted a comparative analysis of degree distributions between our connectomes and three kinds of random graphs: Erdos-Renyi network (Erdos, et al, 1984), small-world network (Watt, et al, 1998), and scale-free network (Barabási, et al, 1999). We utilized NetworkX functions, “gnm_random_graph,” “watts_strogatz_graph,” and “barabasi_albert_graph” for the generation of these random networks, and each kind of random network was generated 100 times independently for a thorough comparison. The parameters employed in the generations were carefully selected to ensure consistency in the number of nodes and approximate number of edges compared to the respective connectomes. The Spearman rank-order correlation coefficient and the p-value (scipy; version 1.21.0) were calculated to estimate the similarity between degree distributions.

For arbor-net and bouton-net, we grouped the cells based on their soma locations, and generated the in-degree and out-degree distributions for each brain region. To compare degree correlation between arbor-net and bouton-net, the degree distributions of brain regions were represented by their median values, and then we calculated the Pearson correlation coefficient (scipy; version 1.21.0) with the medians between arbor-net and bouton-net.

## Supplementary Figures

**Supplementary Figure 1.**
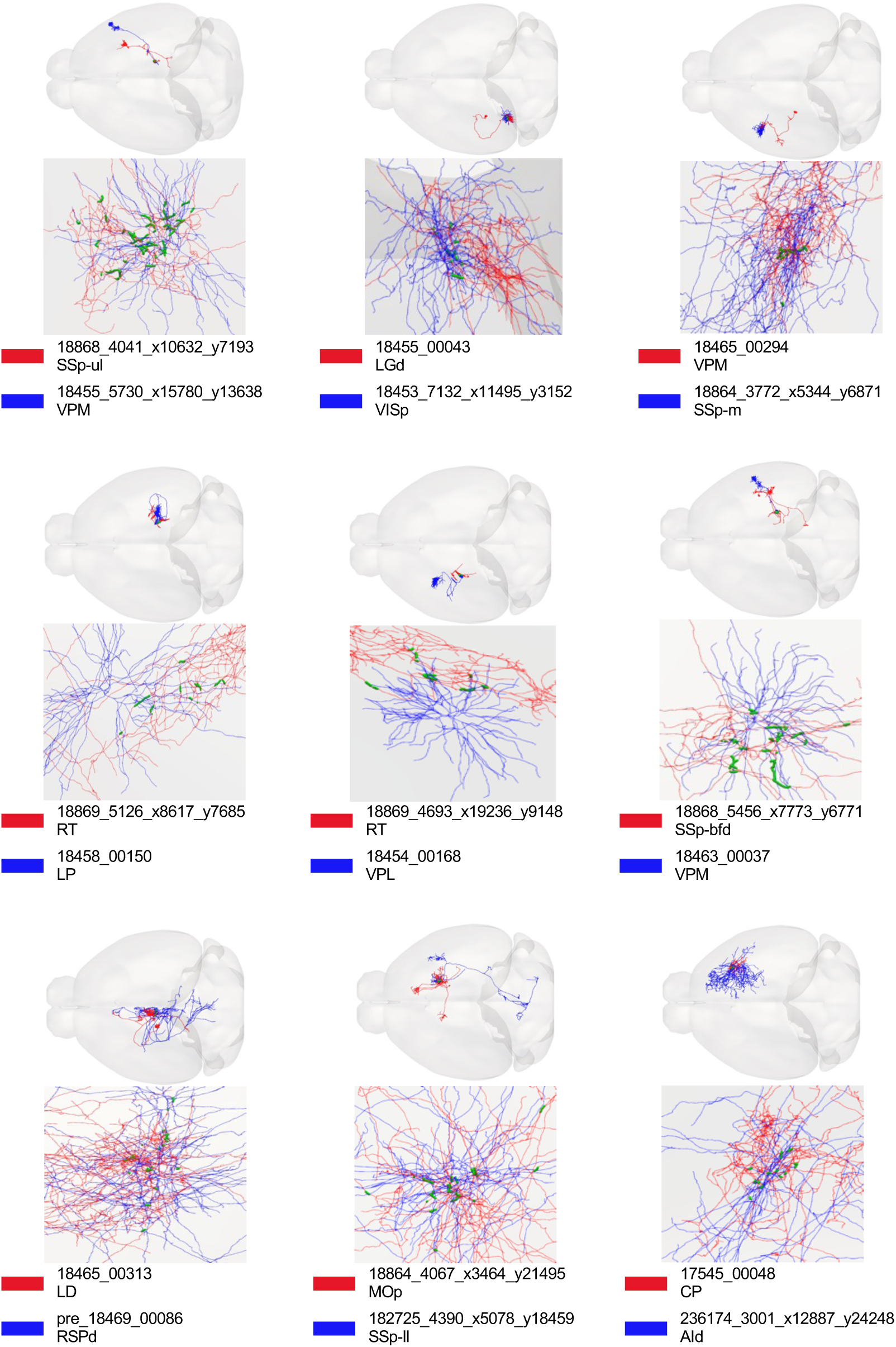
Arbor-net examples. Potential neuronal connections of arbor-net. Colors indicate the source neurons (red) and target neurons (blue). Green dots represent the PACs from these connections.

**Supplementary Figure 2.**
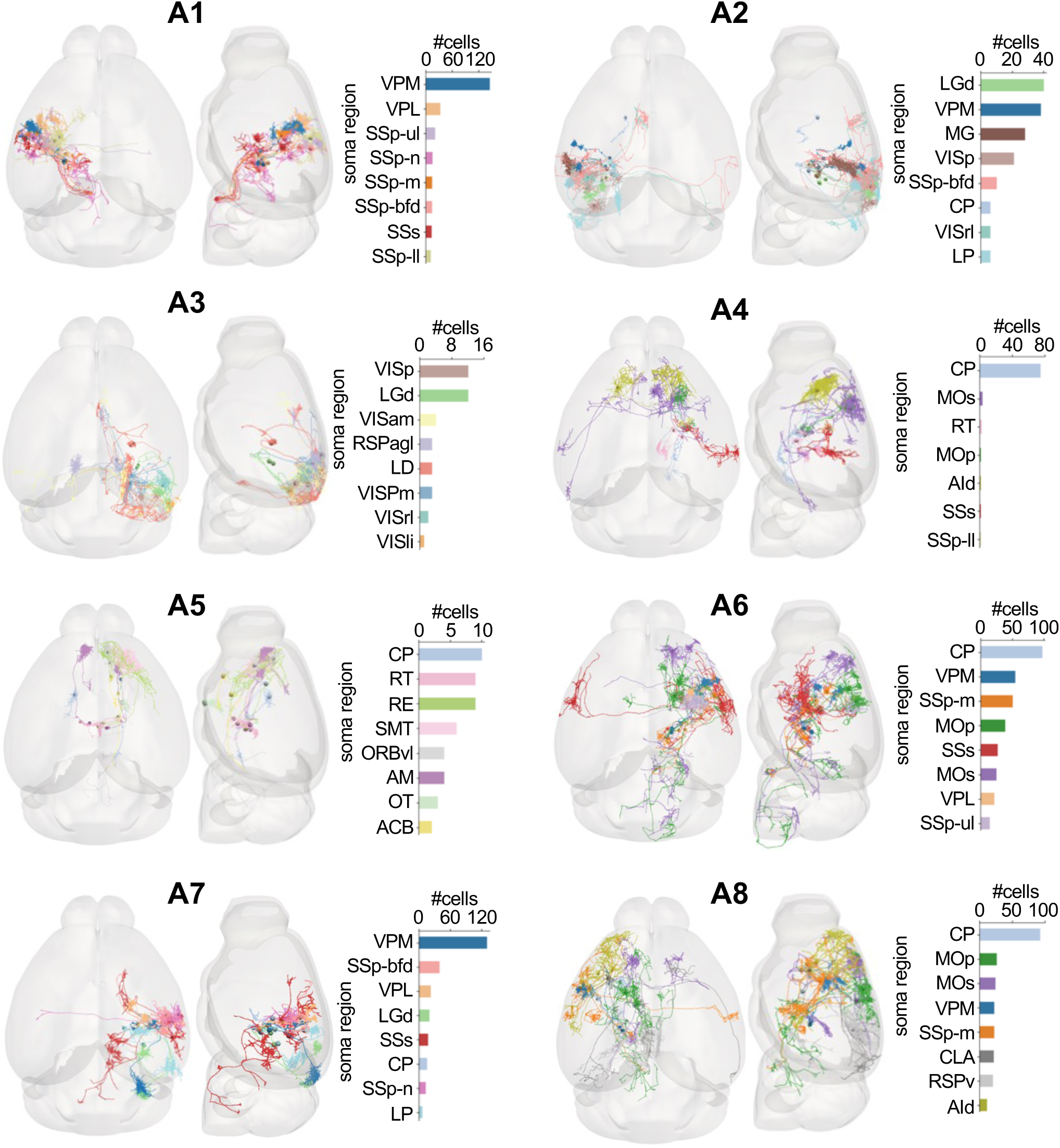
Arbor-net modules. Neuron reconstructions from arbor-net modules are visualized in CCFv3. Colors of neurons correspond to their soma locations. Bar-charts (right): number of cells from main regions for current module.

**Supplementary Figure 3.**
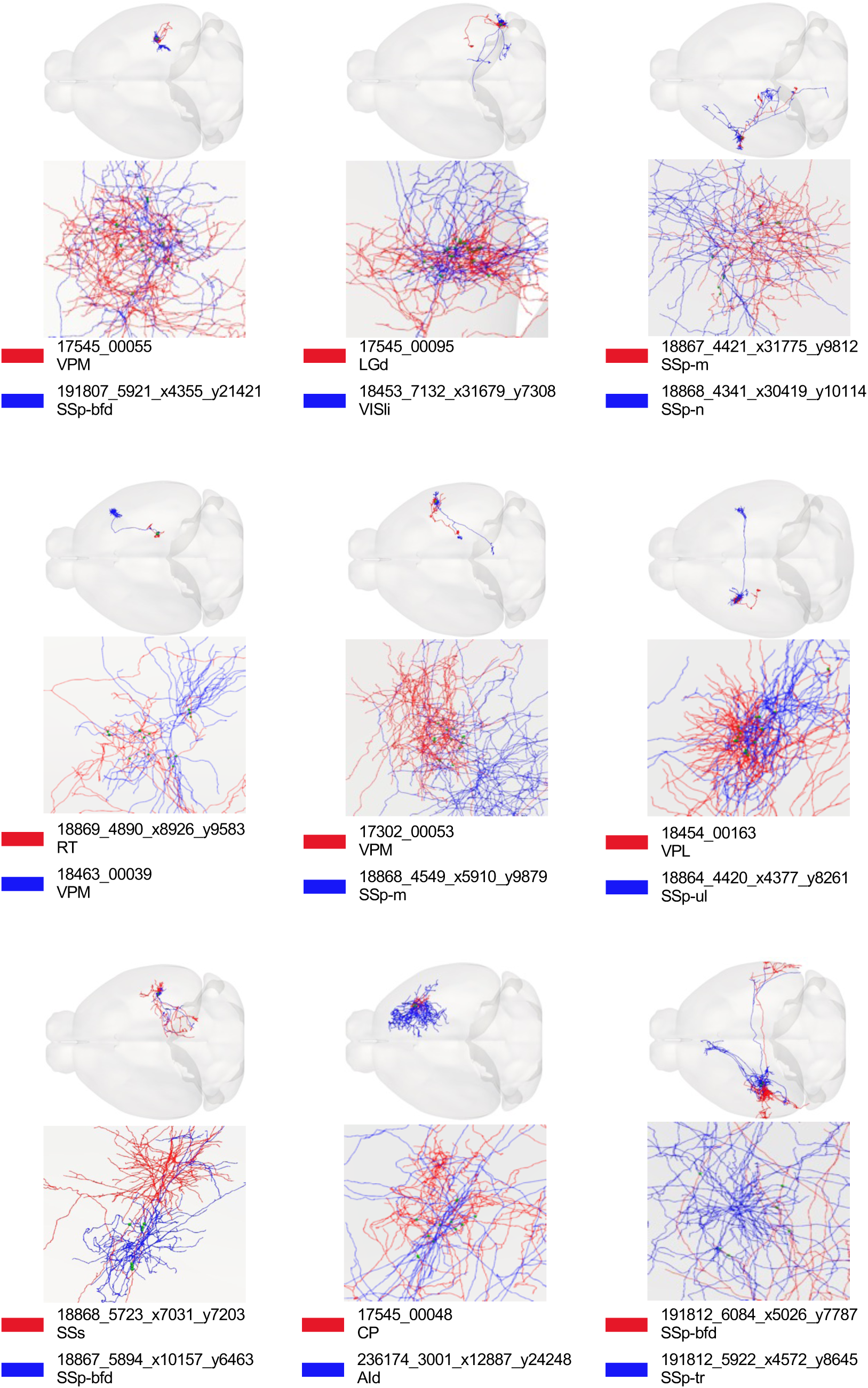
Bouton-net examples. Potential neuronal connections of bouton-net. Colors indicate the source neurons (red) and target neurons (blue). Green dots represent the PBs from these connections.

**Supplementary Figure 4.**
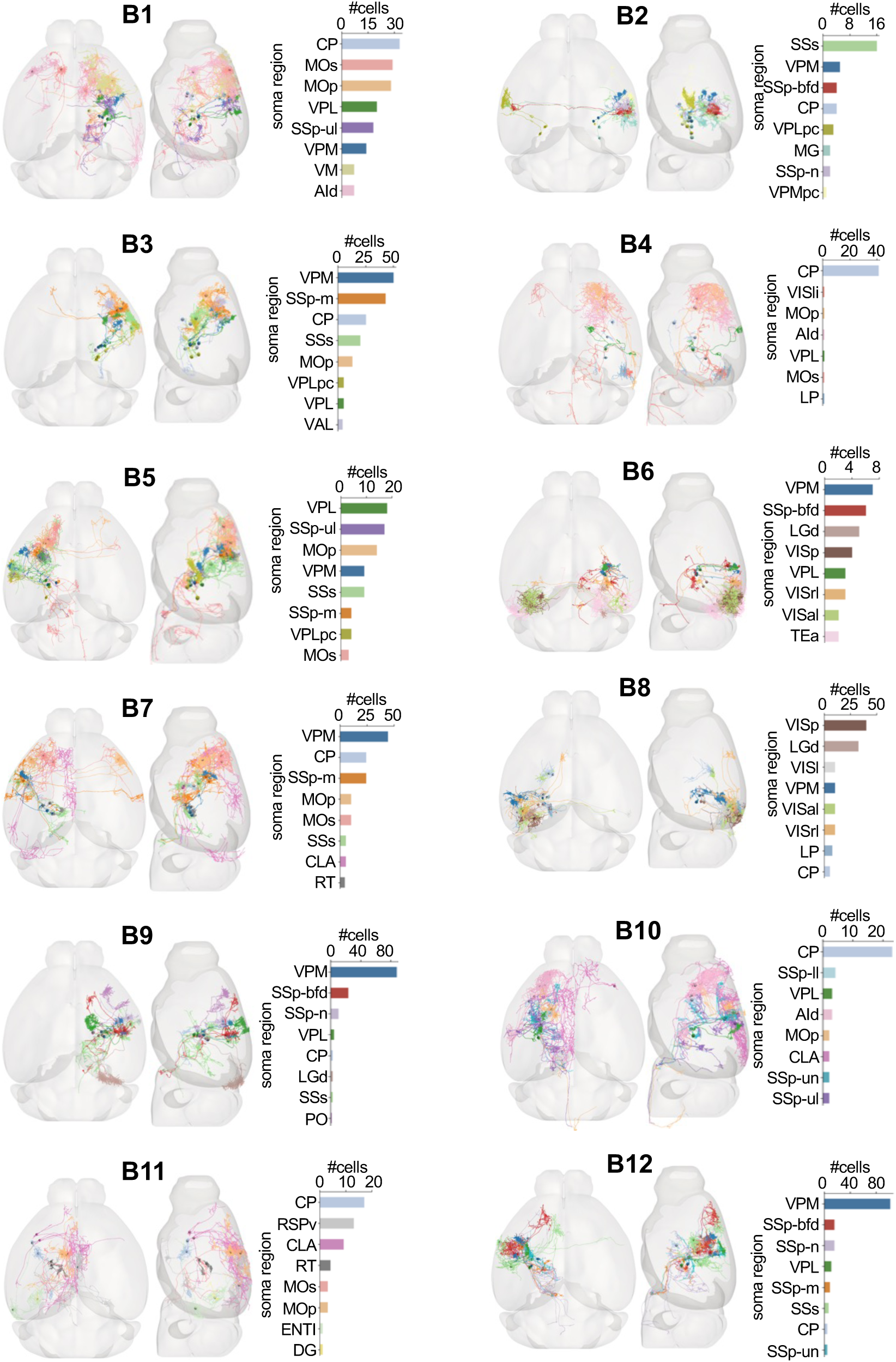
Bouton-net modules. Neuron reconstructions from bouton-net modules are visualized in CCFv3. Colors of neurons correspond to their soma locations. Bar-charts (right): number of cells from main regions for current module.

**Supplementary Figure 5.**
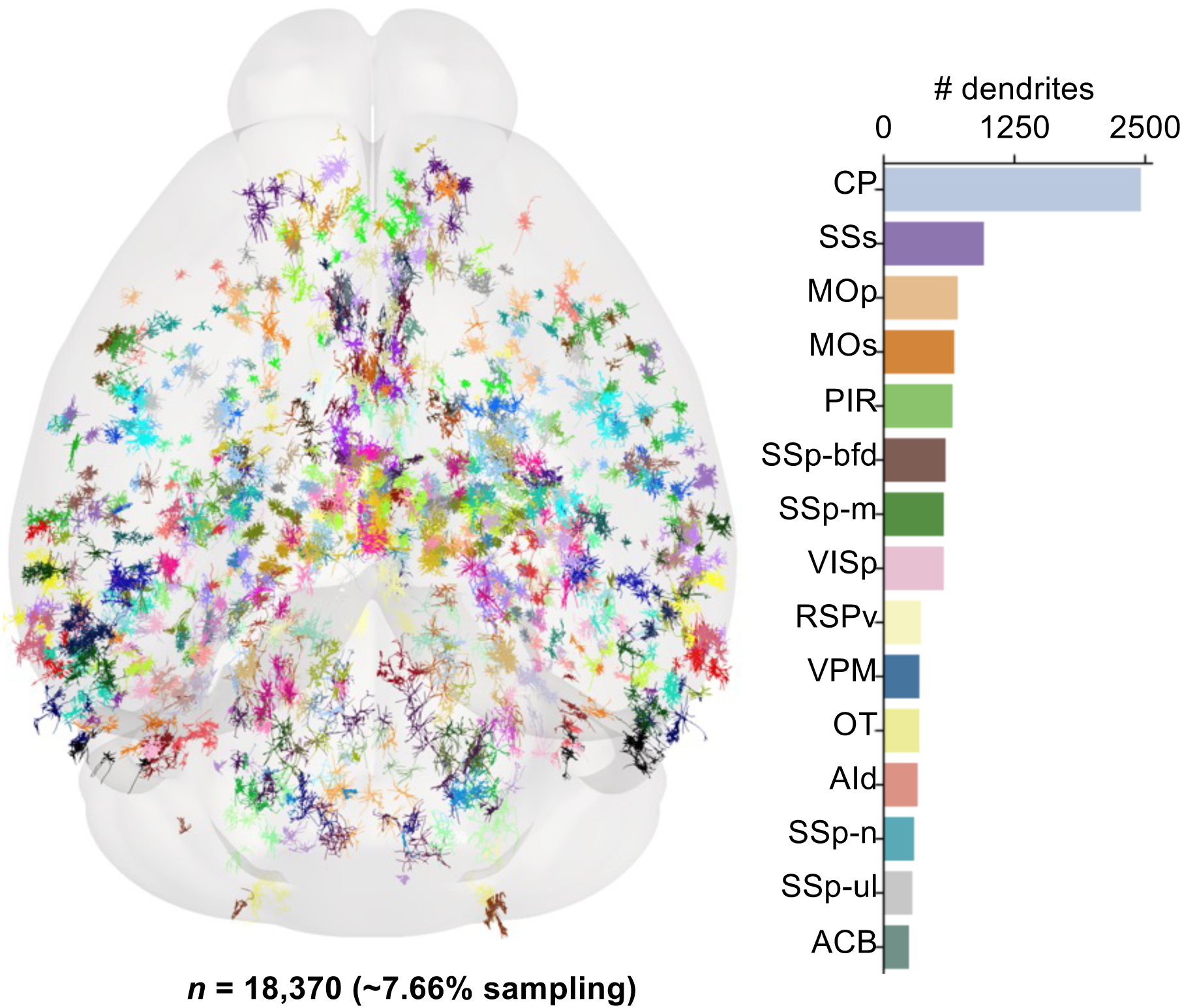
D18370 dendrite examples. Left: dendritic reconstructions from D18370 dataset. Right: number of dendrites in main brain regions.

**Supplementary Figure 6.**
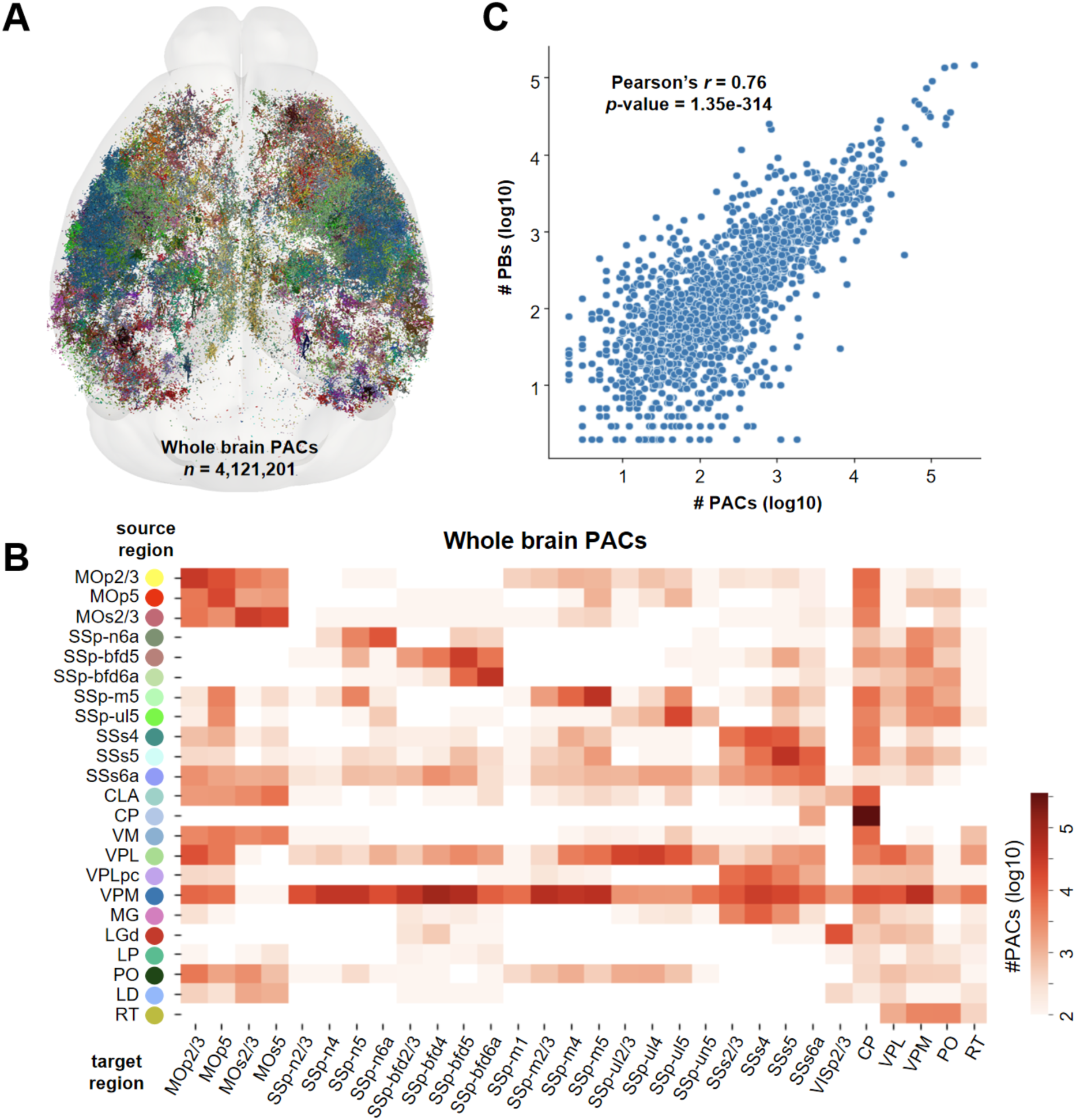
PACs distribution drawn based on D18370 dendrites and F1877 axons. **A.** Spatial distribution of Potential Arbor Contacts (PACs) in whole mouse brain. Colors scheme is same Fig. 4A, indicating their soma locations. **B.** Heatmap of anatomical PACs distribution in whole mouse brain. Horizontal axis: target brain regions of PACs. Vertical axis: source brain regions of PACs. Color bar: the number of PACs (log10-transformed). **C.** Correlation between anatomical distribution of PACs and axonal projection across whole mouse brain. Scatter plot: the counts of PACs and PBs from the same regions.

**Supplementary Figure 7.**
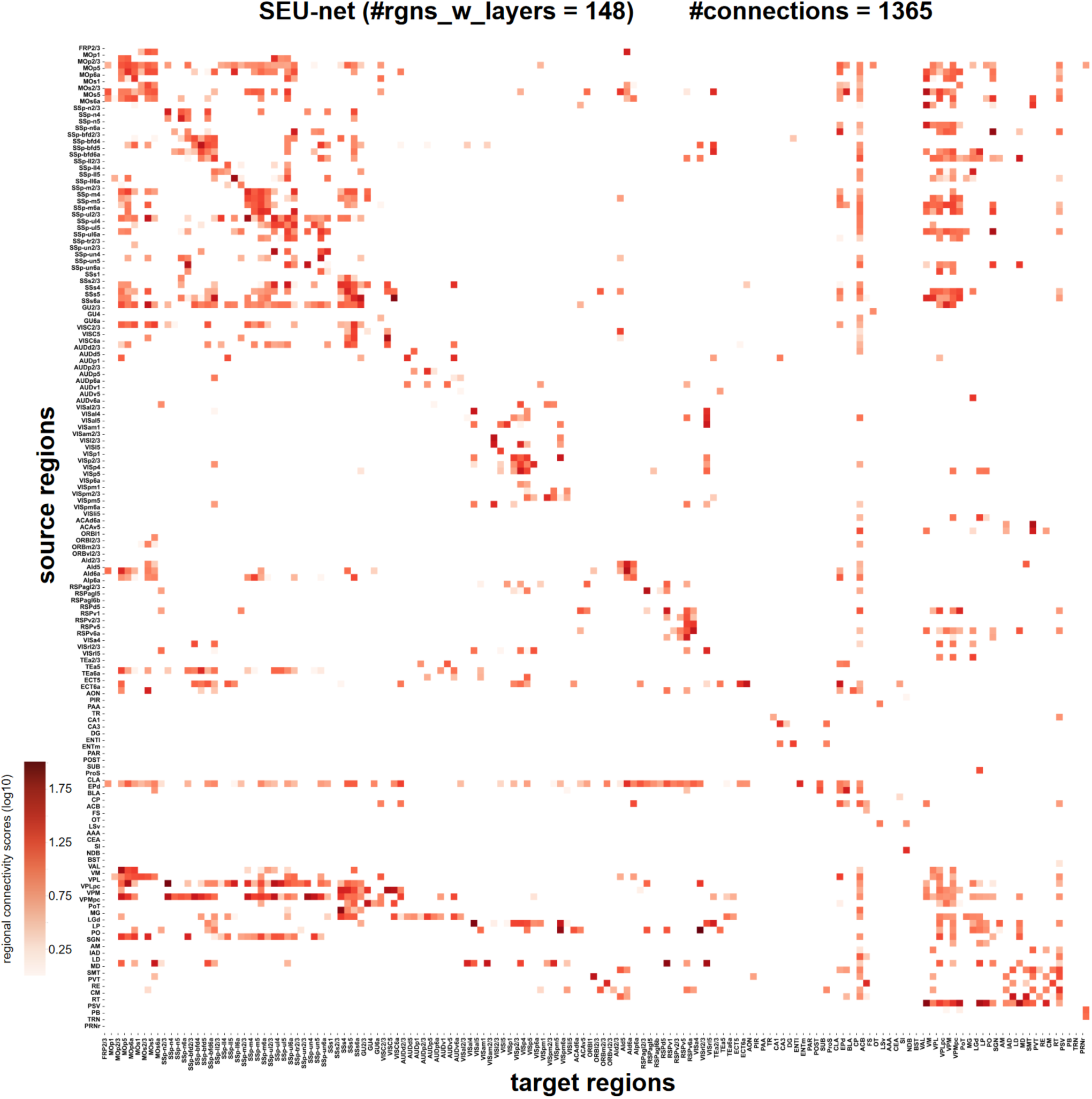
SEU-net: a mesoscale connectome with 148 brain regions including cortical layers. Vertical axis: source brain regions. Horizontal axis: target brain regions. Color bar: regional connection strength (log10-transformed).

**Supplementary Figure 8.**
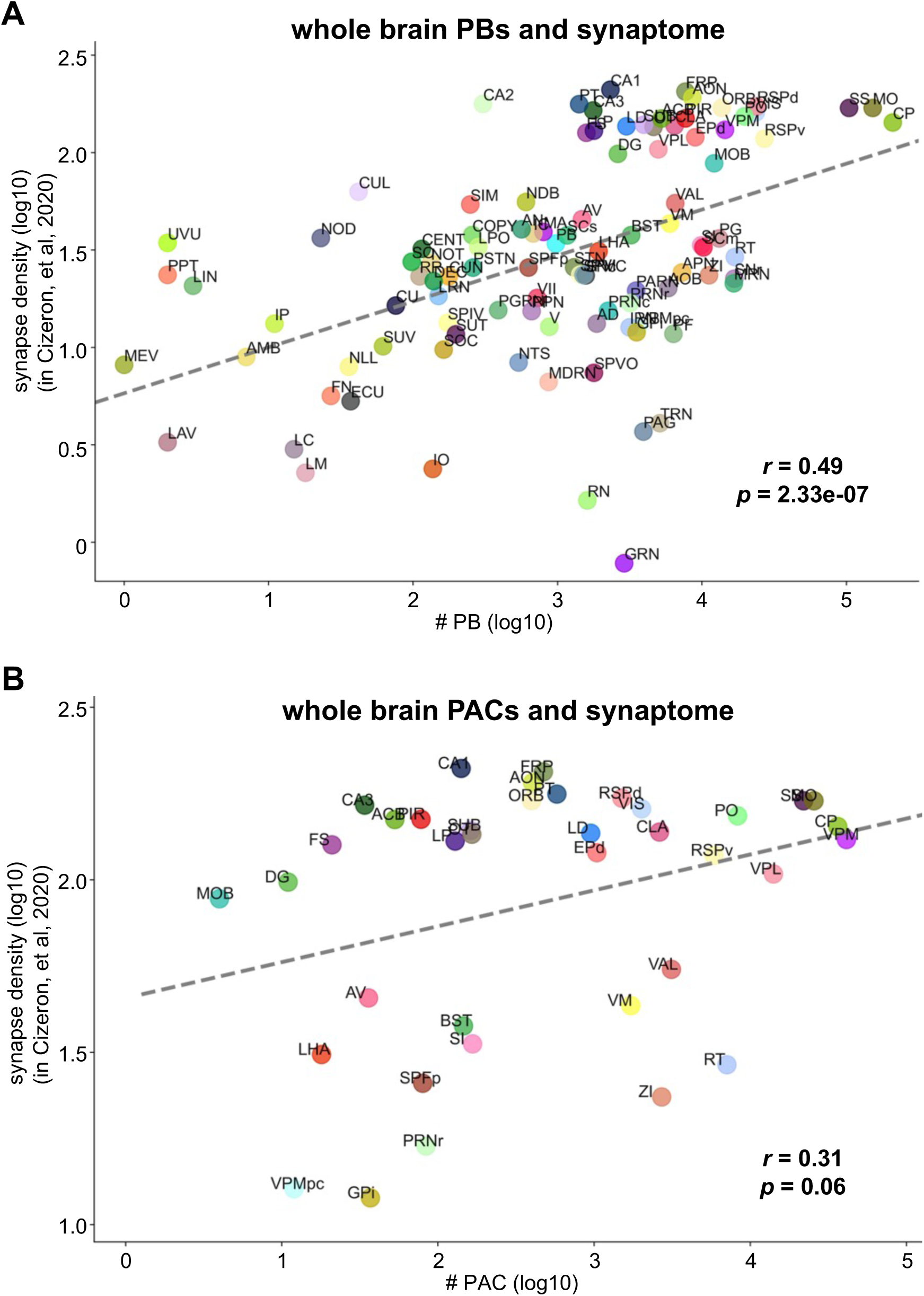
Regional correlation PBs and PACs and a synaptome-based relative density of synapses (Cizeron, et al, 2020). **A.** Correlation between synapse density and anatomical distributions of PBs with 102 shared brain regions. Colored dots: 102 common brain regions. **B.** Correlation between synapse density and anatomical distributions of PACs with 38 shared brain regions. Colored dots: 38 common brain regions. See **Supplementary Table 6** for the list of CCF region names showing up in **A** and **B**.

**Supplementary Figure 9.**
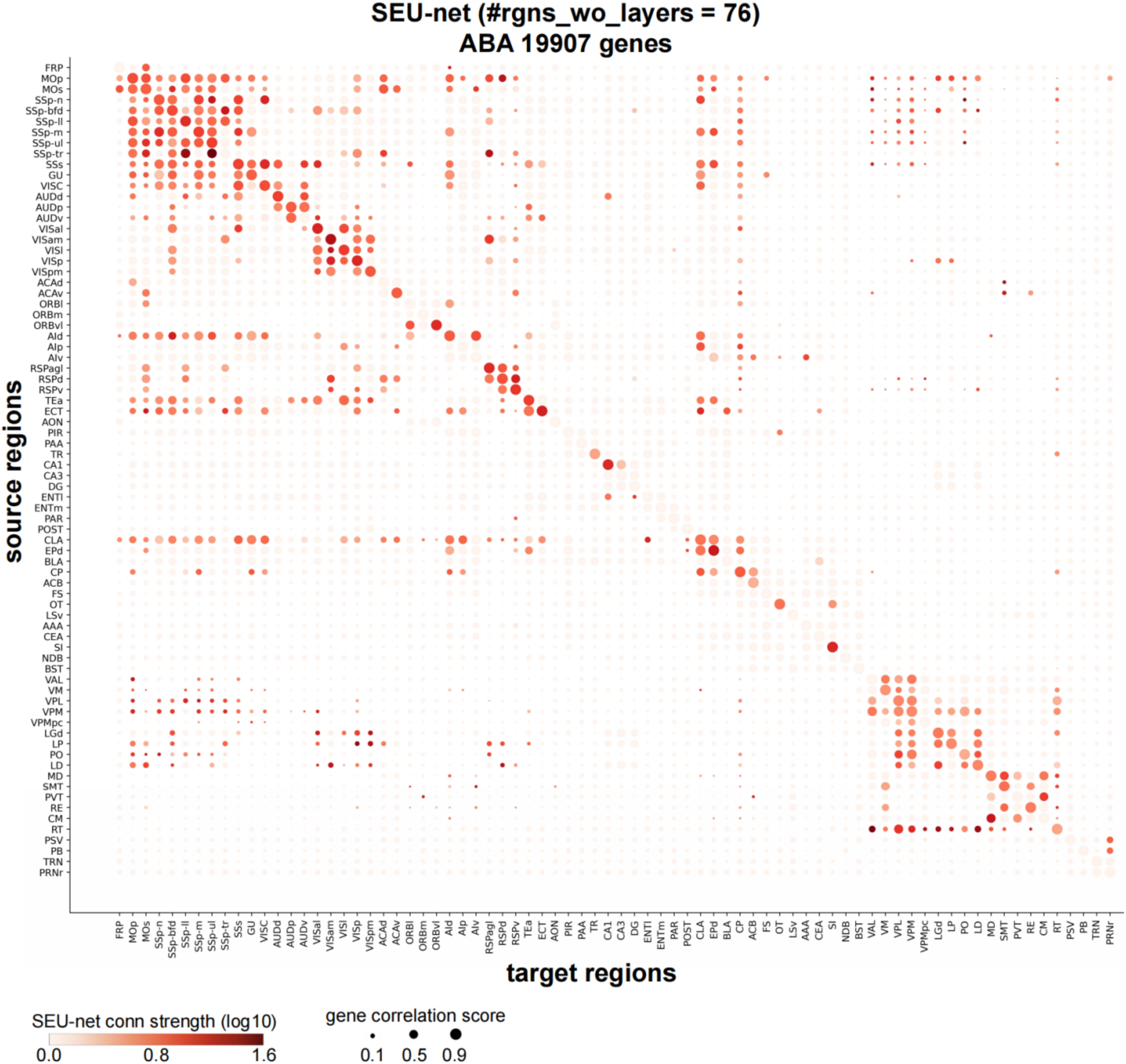
Whole-brain region-wise gene co-expression map using the ABA dataset. Vertical axis: 76 source brain regions. Horizontal axis: 76 target brain regions. Dot colors: regional connectivity of SEU-net. Color bar: regional connection strength from SEU-net. Dot sizes: gene co-expression scores.

**Supplementary Figure 10.**
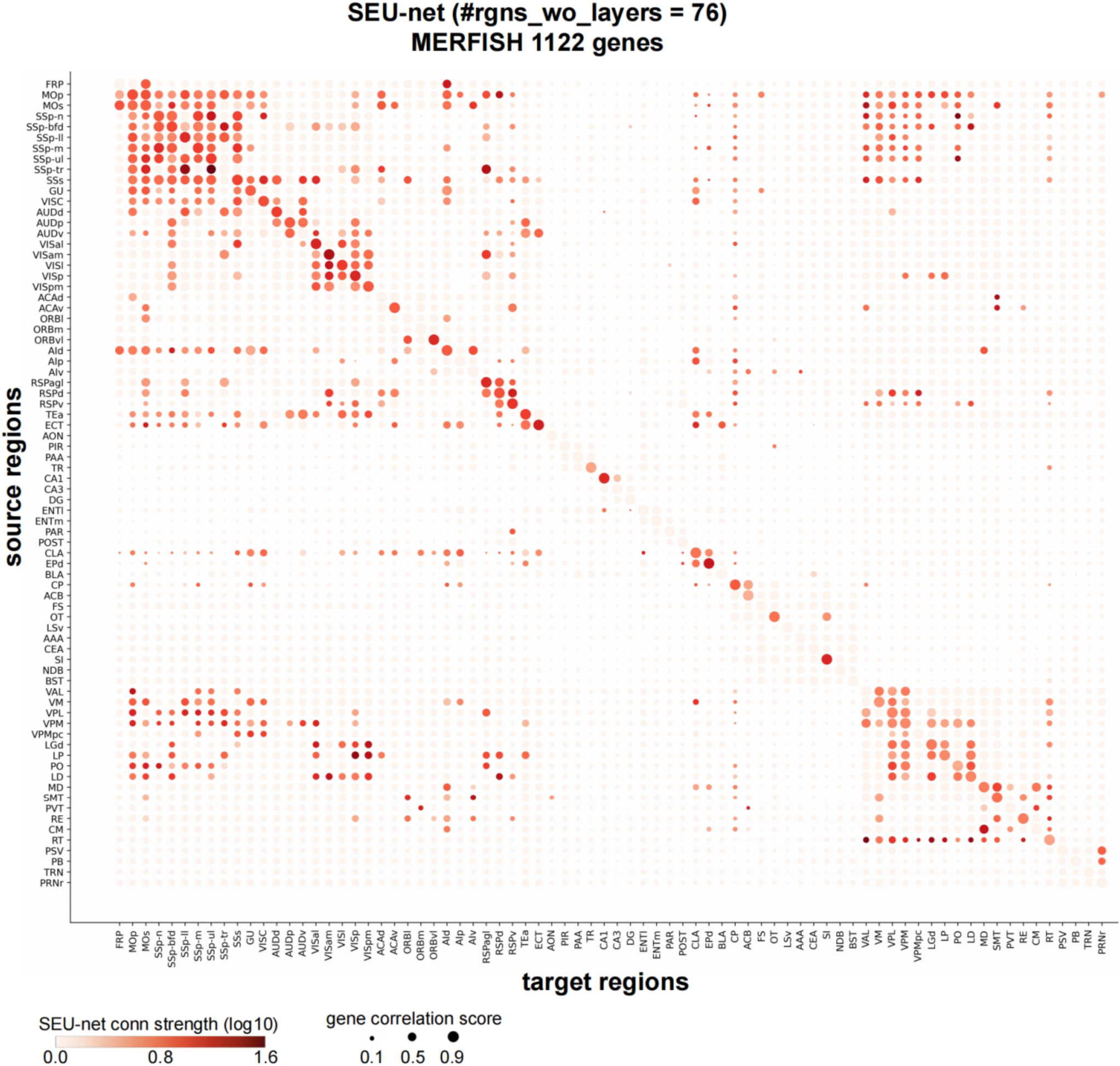
Whole-brain region-wise gene co-expression map using a MERFISH dataset. Vertical axis: 76 source brain regions. Horizontal axis: 76 target brain regions. Dot colors: regional connectivity of SEU-net. Color bar: regional connection strength from SEU-net. Dot sizes: gene co-expression scores.

**Supplementary Figure 11.**
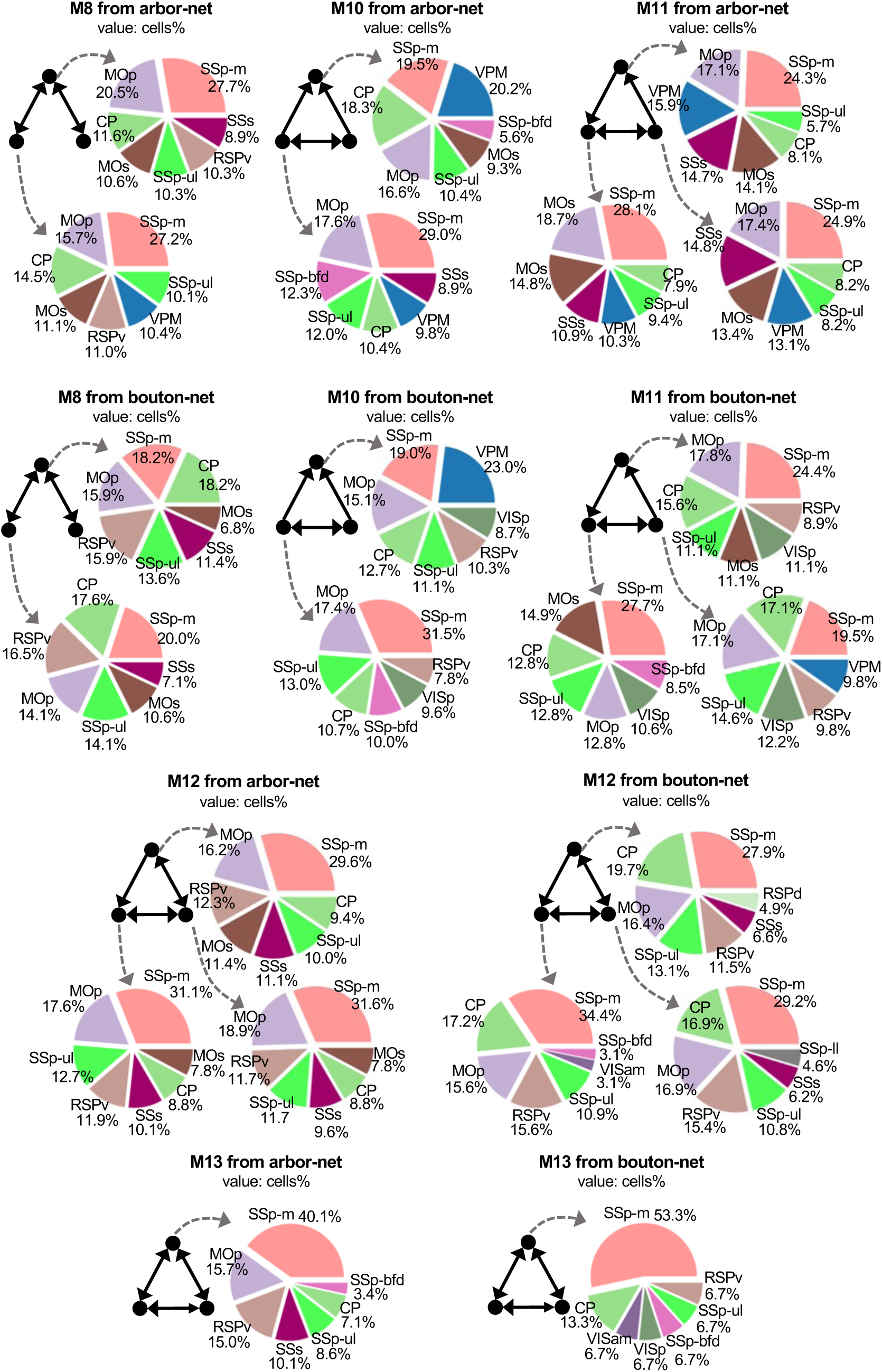
The triads of single neuron connectomes in addition to the M9 motifs shown in Figure 6B.

